# Single-cell transcriptome landscape of circulating CD4^+^ T cell populations in human autoimmune diseases

**DOI:** 10.1101/2023.05.09.540089

**Authors:** Yoshiaki Yasumizu, Daiki Takeuchi, Reo Morimoto, Yusuke Takeshima, Tatsusada Okuno, Makoto Kinoshita, Takayoshi Morita, Yasuhiro Kato, Min Wang, Daisuke Motooka, Daisuke Okuzaki, Yamami Nakamura, Norihisa Mikami, Masaya Arai, Xuan Zhang, Atsushi Kumanogoh, Hideki Mochizuki, Naganari Ohkura, Shimon Sakaguchi

**Author notes:** Correspondence, **Contact info**, Shimon Sakaguchi, Naganari Ohkura. These authors contributed equally to this work (Yoshiaki Yasumizu, Daiki Takeuchi, Reo Morimoto).

## Abstract

CD4^+^ T cells are a key mediator of various autoimmune diseases; however, how they contribute to disease development remains obscure primarily because of their cellular heterogeneity. Here, we evaluated CD4^+^ T cell subpopulations by decomposition-based transcriptome characterization together with canonical clustering strategies. This approach identified 12 independent transcriptional gene programs governing whole CD4^+^ T cell heterogeneity, which can explain the ambiguity of canonical clustering. In addition, we performed a meta-analysis using public single-cell data sets of over 1.8M peripheral CD4^+^ T cells from 953 individuals by projecting cells onto the reference and cataloged cell frequency and qualitative alterations of the populations in 20 diseases. The analyses revealed that the 12 transcriptional programs were useful in characterizing each autoimmune disease and predicting its clinical status. Moreover, genetic variants associated with autoimmune diseases showed disease-specific enrichment within the 12 gene programs. The results collectively provide a landscape of single-cell transcriptomes of CD4^+^ T cell subpopulations involved in autoimmune disease.

## Introduction

Numerous studies have shown that CD4^+^ T cells contribute to autoimmune diseases ^1, 2^, which affect 3-5% of the population and are multifactorial and polygenic ^1, 3^. CD4^+^ T cells exhibit a variety of states (e.g., naive, memory), polarizations (e.g., Th1, Th2, Th17, T follicular helper (Tfh)), and also include a distinct subpopulation engaged in the maintenance of self-tolerance and homeostasis (regulatory T cells (Tregs)) ^4, 5^. While a great deal of effort has been devoted to the detailed classification of CD4^+^ T cells, the complete picture of heterogeneity and its relationship to diseases is still controversial. Furthermore, making consistent assessments across reports is challenging since these reports were based on inconsistent cellular classifications.

The recent emergence of single-cell analysis has greatly contributed to the elucidation of cellular diversities through unbiased profiling ^6–11^. In addition, single-cell RNA-seq (scRNA-seq) is suitable for robust cross-dataset data integration, allowing large-scale investigations ^12–15^. On the other hand, conventional clustering and marker gene detection strategies for single-cell analysis possess the following weaknesses: 1. Cell fraction definition requires arbitrary boundaries; 2. Marker genes for clusters can be occupied by redundant genes or uninterpretable genes, such as long noncoding or ribosomal genes, due to the influence of larger cell population structures; 3. Pairwise differentially expressed gene detection cannot capture global gene variation across multiple clusters. Though some studies have attempted to tackle these issues ^16, 17^, these difficulties have still hindered the interpretations of complex and poorly demarcated cell populations.

Here, we constructed a consensus reference for CD4^+^ T cells in peripheral blood from autoimmune and healthy individuals covering various inflammatory conditions. The reference consists of 18 cell types defined by a conventional clustering strategy and 12 transcriptomic gene programs extracted by conducting decomposition using non-negative matrix factorization (NMF) ^18^ without boundaries, which overcame the weakness of existing single-cell analyses. The results showed that diverse CD4^+^ T cell features were formed by a combination of 12 independent gene programs. We also illustrated that the gene features obtained by NMF could be projected to other bulk / single-cell RNA-seq data to help interpret various datasets. Using these frameworks to examine the genetic contribution and subsequent changes of CD4^+^ T cells in autoimmunity, we performed a meta-analysis that enrolled over 1.8 million CD4^+^ T cells using published single-cell data of 20 diseases and integrated genome-wide association study (GWAS) statistics for 180 traits with our dataset. These analyses provided a full picture of CD4^+^ T cells in autoimmune diseases from the perspective of phenotypes and genetics.

## Results

### Single-cell profiling of peripheral CD4^+^ T cells from healthy and autoimmune donors

To characterize CD4^+^ T cells in various autoimmune properties, we performed single-cell RNA-seq and T cell receptor (TCR)-seq using droplet-based single-cell isolation technology and profiled CD4^+^ T cells, which were collected from three healthy donors, three myasthenia gravis (MG) patients, four multiple sclerosis (MS) patients, and three systemic lupus erythematosus (SLE) patients (Figure 1A; Table S1). After quality control (QC), 103,153 cells were retained and used for the downstream analyses. As the primary layer of clustering (cluster L1), we identified a dynamic differentiation from a naive state via an effector state to a terminally differentiated state. In cluster L1, CD4^+^ naive T cells (Tnaive; *CCR7*^+^ *FAS*^−^), CD4^+^ central memory T cells (Tcm; *CCR7*^+^ *FAS*^+^), CD4^+^ effector memory T cells (Tem; *CCR7*^-^ *FAS*^+^), and CD4^+^ terminally differentiated effector memory T cells (Temra; *FAS*^+^ *CD28*^−^) were observed with distinct gene expression patterns (Figures 1B,D, S1A; Table S2). Tregs were also observed as a distinct cluster with the expression of the master regulator *FOXP3*. Next, we further divided the cells into 18 clusters as the secondary layer, cluster L2 (Figures 1C, S1B-D; Table S3). For example, we broke down cluster L1 cells into several T cell subclusters according to well-known transcription factors and chemokine receptors such as Tcm cells into Tfh (Tfh; *CXCR5*, *PDCD1*), Th2 (*GATA3*, *CCR4*), Th17 (*RORC*, *CCR6*); Tem cells into Th1/17 (*TBX21*/Tbet, *RORC*), Th1 (*TBX21*/Tbet); Temra cells into Th1 (Figures 1C,D, S1A-D). Treg cells were divided into three clusters; Treg Naive (*CCR7*), Treg Activated (*ID2*), and Treg Effector (*CCR4*) (Figures 1C,D, S1A-D). In addition, several minor clusters were found, such as Tnaive *MX1*, which preferentially expresses interferon signature genes (Figures 1C,D, S1A,C,D). Transcriptome profiles of each cluster were concordant with bulk RNA-seq data from sorted CD4^+^ T cell fractions provided by the DICE consortium ^19^ (Figure 1E). We found a *CXCR5*^-^ *PDCD1*^+^ cluster occupying 1% in CD4^+^ T cells whose marker genes corresponded to the canonical marker for T peripheral helper (Tph) cells ^20–22^ in Tem (Figures S1A,E). The population was annotated as circulating Tph, although a few cells with the expression *CXCR5*^-^ *PDCD1*^+^ were also observed in broader populations, such as Tcm and Temra, (Figures S1A,C,F). Overall, we identified cell populations of peripheral CD4^+^ T cells from healthy and autoimmune states using scRNA-seq.

**Figure 1.**
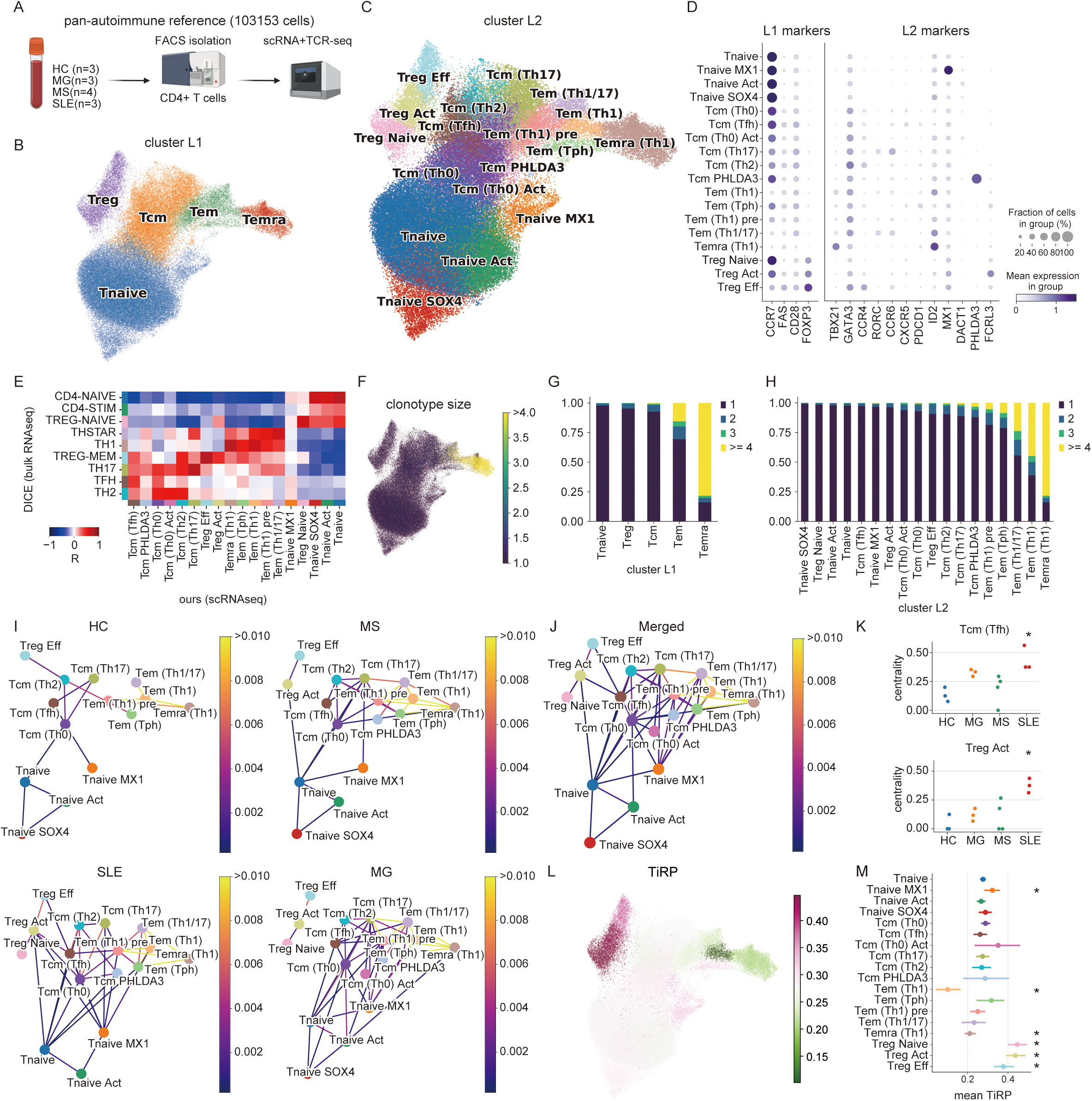
Global profiling of CD4^+^ T cells. (A) Sample collection strategy. (B and C) Clusters L1 and L2 on UMAP embeddings. (D) Dot plot depicting signature genes’ mean expression levels and percentage of cells expressing them across clusters. Marker genes for the plot were manually selected. See also Figure S1C for automatically extracted marker genes. (E) Expression correlation of clusters with bulk RNA-seq for sorted CD4^+^ T cell fractions (DICE). (F) UMAP plot showing clonotype size. (G and H) Clonotype size distributions across clusters. (I and J) TCR similarity networks in autoimmune patients, healthy donors (I), and all donors (J). TCR similarity was calculated for each sample, and only edges where overlapping clonotypes were detected in >=2 (I) or >=4 (J) samples are depicted as robust overlaps. The edge color indicates the average TCR similarity of all samples. (K) Degree centrality of TCR networks. Significance across clusters was calculated by one-way ANOVA, and after multiple test corrections by FDR, Tcm (Tfh) and Treg Act were retained as significant cell types. Then, pairwise Tukey-HSD posthoc tests were performed. *: p_adj_ < 0.05 in comparison with HC. (L) TiRP score distributions on UMAP plot. Mean scores for each cluster L2 were shown. (M) TiRP score distribution across cluster L2. The dot shows the mean, and the CI shows 95% CI of the bootstrap distribution of means (n=1000). Adjusted p-values of significant clusters, Tnaive *MX1* 1.29×10^-2^, Tem (Th1) 5.82 x 10^-11^, Temra (Th1) 4.84×10^-22^, Treg Naive 2.14 x 10^-13^, Treg Act 2.14 x 10^-13^, Treg Eff 3.82 x 10^-5^ (Two-sided Mann-Whitney’s U-test was performed for one cluster vs. the other clusters iteratively).

### TCR features across CD4^+^ T cells reflect cellular properties

Because TCR responses shape T-cell functions and differentiation, TCR diversities and overlaps provide useful information for the properties and relationships of populations. Therefore, we analyzed single-cell TCR features sequenced along with gene expression. Clonotype sizes and diversity across cluster L1 populations revealed that Temra was most clonally expanded, followed by Tem, Tcm, Treg, and Tnaive (Figures 1F,G). Similarly, in cluster L2 populations, Temra (Th1), Tem (Th1), and Tem (Th1/17) possessed a limited number of clonotypes, whereas Tnaive and Treg Naive maintained diverse clonotype pools (Figure 1H). TCR similarity network showed repertoire sharing between neighboring clusters, Tnaive and Tcm, Tcm and Tem, Tem and Temra, while the distal connection, such as from Tnaive to Temra was not observed, suggesting stepwise development from Tnaive to Tcm, Tem, and Temra (Figures 1I,J). The repertoires were also mutually shared within Tcm cell populations, suggesting the plasticity of T cell polarization against the same epitopes. In addition, Treg Naive and naive conventional T cells (Tconvs) didn’t share repertories, whereas Treg Act and Treg Eff shared repertoires with Tcm populations. We also measured the centrality of TCR networks for each cell type to evaluate the differentiation potential of each cluster. The centrality of Tcm (Th0), Tem (Th1) pre, and Tnaive were consistently high, suggesting that these cells possess the possibility to differentiate into a variety of cell types (Figure S2A). In addition, TCR networks differed depending on the disease states (Figure 1I). Especially the centrality of Tcm (Tfh) and Treg Act were higher in SLE (Figures 1K, S2B). These results indicated that the kinetics of CD4^+^ T cell differentiation varied depending on the disease state.

Previous studies have shown that T cells with stronger TCR stimulation within the thymus are more likely to differentiate into Tregs than Tconvs ^23^. Therefore, Tregs have specific TCR properties, such as hydrophobicity in complementary determining region 3 (CDR3) regions ^24^. We measured the Tregness of TCRβ chains (TCR-intrinsic regulatory potential, TiRP score ^24^) and found that the mean TiRP score was higher in Treg cells compared with Tconv cells (Figures 1L,M). On the contrary, Tem (Th1) showed a low TiRP score indicating that Tem (Th1) has experienced the stimulation with non-self antigens. Among Treg cells, Treg Naive and Treg Act showed higher TiRP scores than Treg Eff. It has been thought that naive Tregs contain predominantly thymic differentiated Tregs (tTregs), while effector Tregs are compensated by peripherally differentiated Tregs in addition to tTregs ^25^. This notion was concordant with our observations that naive Tregs had the strongest Treg characteristics in the TCRs and that Treg Act and Eff shared TCRs with Tconvs (Figures 1I,J, S2C). Furthermore, in MS patients, the TiRP scores of the Treg Act were significantly low, reflecting disease-dependent Treg compensation by Tconvs (Figure S2D). Overall, TCR repertoires provided valuable insights into T cell characteristics and relationships during the differentiation.

### Decomposition of cellular programs using NMF

Next, we attempted to identify cellular programs within and across cell types. We noticed that conventional clustering and marker gene detections could fail to capture meaningful clusters and genes. For example, differentially expressed genes in our reference included overlapping genes among Th1 cell populations and nonsense genes in Tnaive cells, suggesting the conventional marker gene detection is insufficient for CD4^+^ T cells (Figure S1C). We suspected that artificially delineating in the clustering process is unsuitable for a gradual population such as CD4^+^ T cells. In addition, because marker gene detections are performed by pairwise comparison, global representations across cell types cannot be detected. To overcome these limitations, we applied non-negative matrix factorization (NMF) ^18^ to normalized gene expression of our scRNA-seq data and unbiasedly dissected gene expression profiles into a gene feature matrix ***W*** and a cell feature matrix ***H*** (Figure 2A). To determine the number of components, we assessed the explained variances and maximum inter-component correlations and selected 12 for the number of components as they kept sufficient information and were not redundant (Figure S3A; methods). Based on the gene feature profiles and the enriched pathways, we annotated the NMF components (Figures 2B-D, S3B; Table S4,5). Several factors were related to T-cell polarization, such as Treg-Feature (Treg-F, NMF1; genes with high weights; *IKZF2*, *FOXP3*), Th17-F (NMF 2; *RORC*, *CCR6*), TregEff/Th2-F (NMF5; HLA class II genes, *CCR10*, *CCR4*), Tfh-F (NMF6; *TIGIT*, *CXCR5*), Th1-F (NMF11; *GZMK, EOMES*, *CXCR3*), and differentiations such as Naive-F (NMF3; *CCR7*, *TCF7*), Central Memory-F (NMF8; *S100A8*, *ANXA1*), and Cytotoxic-F (NMF0; *GZMB*, *NKG7*). NMF5 was enriched in both Th2 and Treg Eff, suggesting that effector Treg cells and Th2 cells may be controlled by the shared program as previously suggested ^26^ (Figure 2B). NMF6 (Tfh-F) also demonstrated moderate activity in Treg Act, suggesting an overlap between Treg Act and T-follicular regulatory (Tfr) cells ^27^ (Figure 2B). NMF11^high^ cells were enriched in Tem (Tph), Tem (Th1), and Tem (Th1/17) cells showing a wide range of Th1ness gene usage across these subtypes. Moreover, NMF7 was a type I interferon signature gene component enriched in Tnaive *MX1* (Figures 1C, S3B). Intriguingly, NMF10 captured a global feature across cell types consisting of AP-1 family genes (*JUNB*, *FOS*), *NFKBIA*, *CD69*, and *CXCR4* (Figure 2D). This feature was concordant with tissue-homing T-cells observed in the thymoma of MG patients ^7^ and the central nervous system of neurodegenerative disease patients ^28^, and was labeled as Tissue-F. NMF4 (Act-F) was related to *IL7R* signaling, which is an essential survival and differentiation signal ^29^. The proportion of explained variance (Evar) showed the most drastic variations in the peripheral CD4^+^ T cells were differentiation from Tnaive to Tcm, Tem, and Temra, and the polarizations were relatively smaller changes and independent of the differentiation programs (Figure 2B). Altogether, NMF succeeded in the decomposition of peripheral CD4*^+^* T cell gene programs into 12 components and showed that complex CD4*^+^* T cell populations were represented by a simple combination of the 12 components.

**Figure 2.**
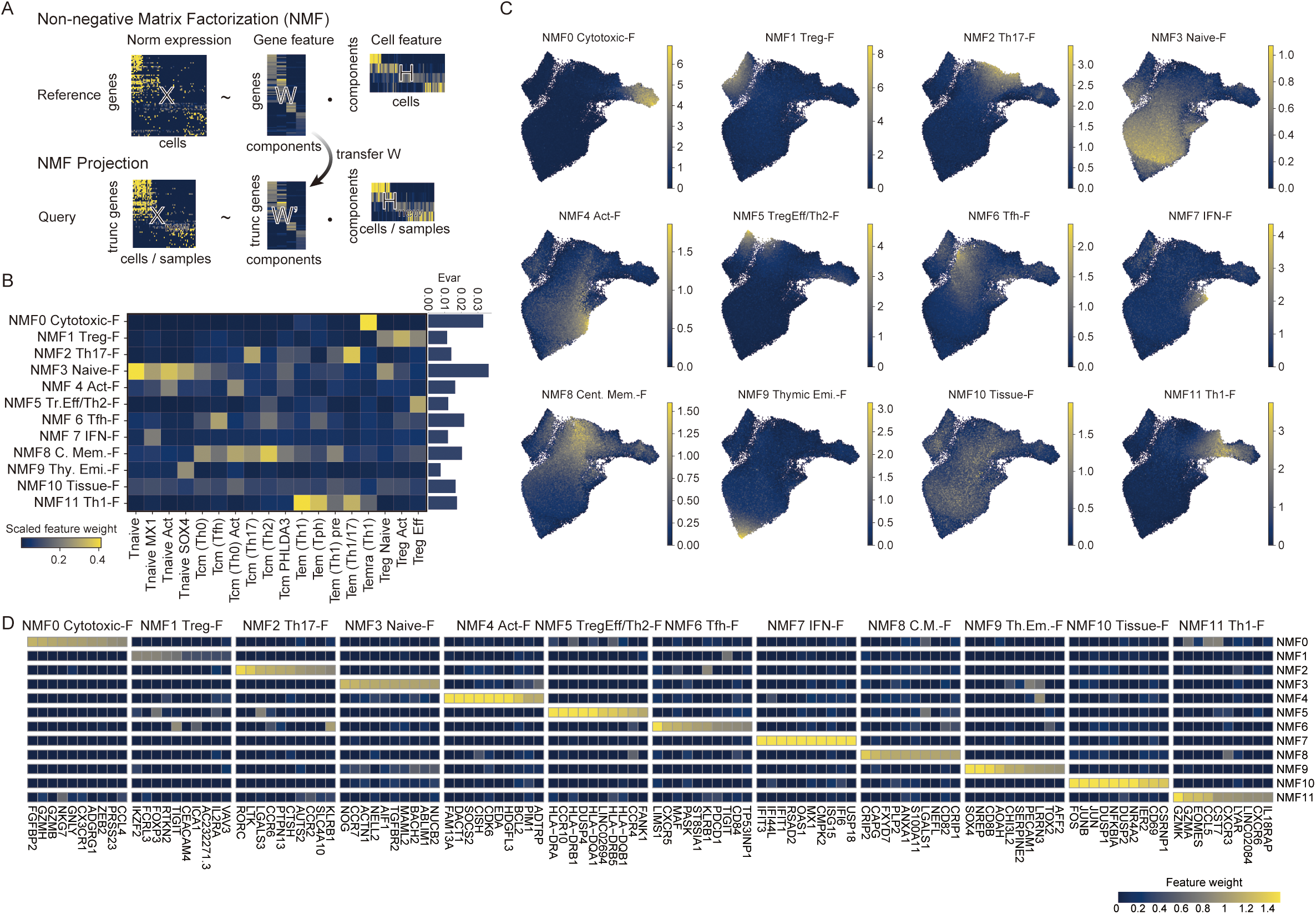
NMF captured 12 CD4^+^ T cell features. (A) Schematic view of NMF and NMF projection. (B) Matrixplot showing the mean scaled NMF feature weight for each cluster L2 population. The explained variance (Evar) is also shown on the right. The NMF feature weight is scaled by the maximum value for each feature for visualization. (C) NMF cell feature value on UMAP plots. (D) Gene features for each component. The top 10 genes for each feature were selected. The 12 gene features are annotated using top genes and previous reports as NMF0 Cytotoxic-Feature (F), NMF1 Treg-F, NMF2 Th17-F, NMF3 Naive-F, NMF4 Activation-F (Act-F), NMF5 TregEff/Th2-F, NMF6 Tfh-F, NMF7 IFN-F, NMF8 Central Memory-F, NMF9 Thymic emigrant-F, NMF10 Tissue-F, NMF11 Th1-F.

### NMF projection enables fast interpretation of various CD4^+^ T transcriptome datasets

One of the biggest challenges in single-cell analysis is the integration of datasets. To achieve a simple integration, we expanded the NMF framework to allow the projection of the pre-computed gene feature matrix onto other datasets by developing a bioinformatics tool, NMFproj (Figure 2A, https://github.com/yyoshiaki/NMFprojection). To measure how the NMF features explain the variance of the query dataset, we introduced a QC metric named the proportion of overlapped highly variable genes (POH) (Figure S3C). A low POH indicates that the query data set has much variability other than the NMF features evaluated by NMFproj. We applied NMFproj to various datasets to validate the scalability (Supplementary Note). Analysis of bulk RNA-seq of sorted peripheral CD4^+^ T cells provided by the DICE project ^19^ demonstrated that each fraction was well represented by the 12 NMF gene features (POH: 0.272, Figure S3D). Miyara classification ^30^, which classified CD4^+^ T cells into Fr. I to Fr. VI by the expression of CD45RA and CD25, was re-evaluated by NMFproj and showed that Fr. III (CD45RA^-^ CD25^int^) has Th17 type characteristics in line with the original report ^30^ (POH: 0.542, Figure S3E, Supplementary Note). In addition, profiling of circulating Tph cells ^20^ revealed that Tph cells possessed both NMF6 (Tfh-F) and NMF11 (Th1-F) in concordance with Tem (Tph) we defined as cluster L2 (POH: 0.134, Figure S3F, Supplementary Note). We also attempted to utilize NMFproj for the QC of *in vitro* induced Treg (iTreg) cells of mouse ^31^ and found that iTreg cells induced in optimized conditions for enhancing Treg functionality showed higher NMF1 (Treg-F) values than conventional iTreg cells (POH: 0.182, Figure S3G, Supplementary Note). We also applied NMFproj to scRNA-seq datasets of cross-tissue immune cells ^32^ (Figure S5, POH: 0.560 in CD4^+^ T cells, Supplementary Note), pan-cancer tumor-infiltrating CD4^+^ T cells ^15^ (Figure S4A, POH: 0.530, Supplementary Note), and mouse splenocytes ^33^ (Figure S4B, POH: 0.394, Supplementary Note), achieving robust interpretations of cellular features in various conditions. Furthermore, cell-specific qualitative changes have been reported in autoimmune diseases, such as Treg dysfunction in SLE ^34^, and we hypothesized that NMFproj could be used to detect these changes in individual cell populations. To test this, we applied NMFproj to bulk RNA-seq data of sorted peripheral CD4^+^ T cell fractions from various autoimmune patients ^35^. NMFproj detected a subset-specific gene program robustly even in a variety of autoimmune disease conditions (Figure S6A, Supplementary Note). The results showed cell-type wide enhancement of NMF7 (IFN-F) in SLE and mixed connective tissue disease (MCTD) and hampered NMF1 (Treg-F) in Fr.I nTregs (CD45RA^+^ CD25^+^) in SLE patients as previously reported ^34^ (Figure S6B). These results indicated that NMFproj could robustly assess the qualities of CD4^+^ T cells in various tissues and disease states, regardless of bulk/single cell or human/mouse.

### Meta-analysis of CD4^+^ T cells in various autoimmune diseases

To extend CD4^+^ T cell profiling to various autoimmune and infectious diseases, we performed a meta-analysis using publicly available single-cell data ^6, 8, 36–57^. We integrated publicly available datasets with two strategies: 1) quantitative evaluation of cell frequencies by mapping to our reference and 2) evaluation of qualitative changes per cell type using NMFproj. We extracted CD4^+^ T cells from peripheral blood mononuclear cells (PBMCs) using Azimuth ^58^ and then mapped them to our reference using Symphony ^14^ (Figure 3A, the pipeline is available at https://github.com/yyoshiaki/screfmapping). We collected 1,809,668 CD4^+^ T cells collected from 647 cases and 306 controls from 25 projects (Figures 3B, S7A; Table S6). For quality assurance, only datasets in which both HC and patients were present and at least 3 cases were included were used. As a prominent change, Tnaive decreased, and Temra increased in various autoimmune diseases (Figure S7B; Table S7). It has been reported that Temra increased in the peripheral blood of rheumatoid arthritis (RA), MS, ulcerative colitis (UC), and Crohn’s disease (CD) patients ^59–62^, which was consistent with the present data. Kawasaki disease and Type 1 diabetes (T1D) were exceptions among autoimmune diseases, with a slight increase in Tnaive and no significant change in Tcm, Tem, and Temra (Figure S7B), as reported previously ^63, 64^. At cluster L2 resolution, we found that Tnaive *MX1* increased in COVID-19, SLE, T1D, and primary Sjögren syndrome (pSS) patients (Figure 3C). The type I IFN response is essential for viral elimination and has been reported to be associated with COVID-19 pathology ^65^ and also known to be associated with SLE ^66^, pSS ^67^, and T1D ^68^.

**Figure 3.**
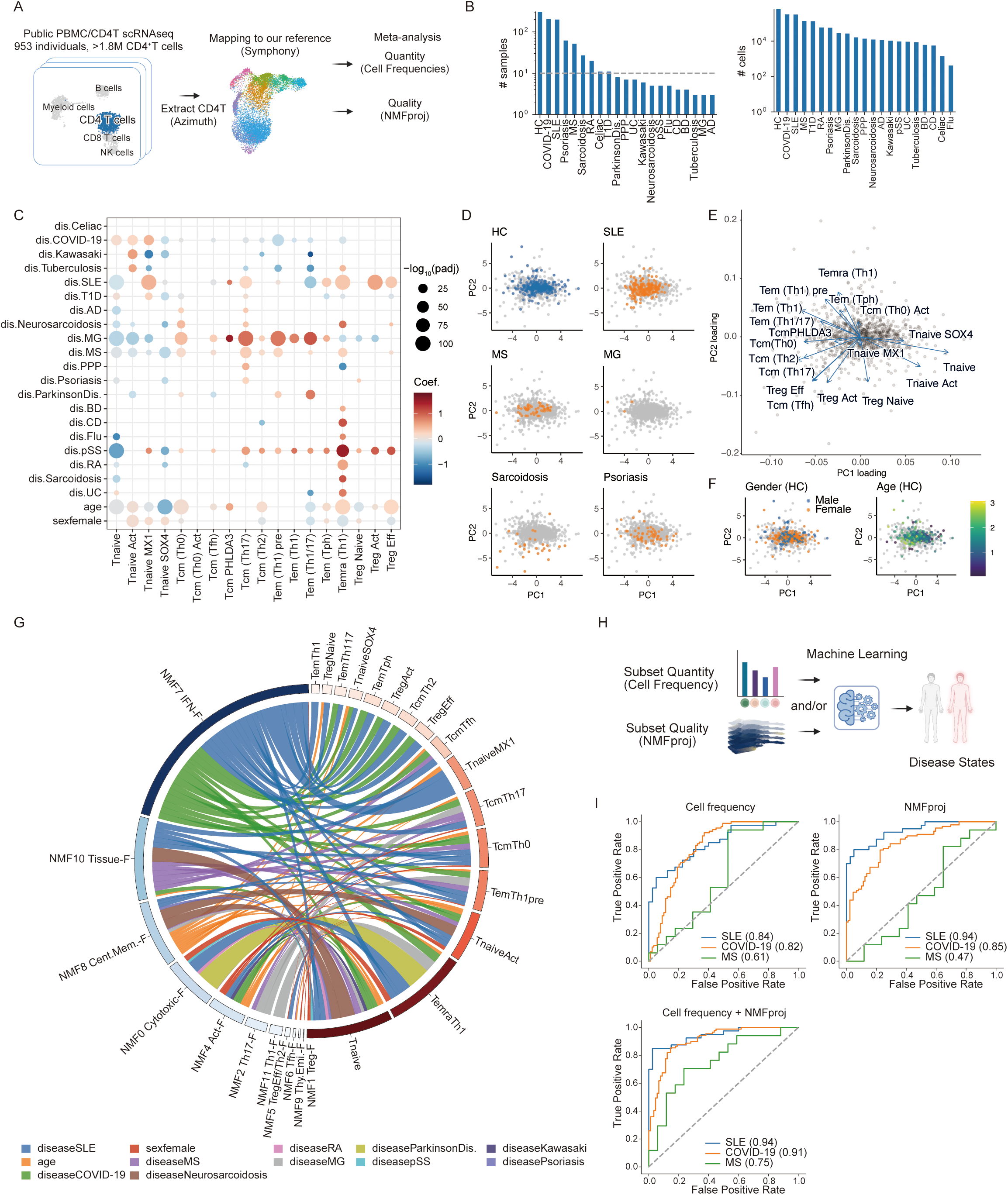
Pan-autoimmune meta-analysis of peripheral CD4^+^ T cells. (A) Strategy for meta-analysis of peripheral CD4^+^ T cells across diseases. First, CD4^+^ T cells were extracted from PBMC scRNA-seq datasets using Azimuth ^58^. Extracted CD4^+^ T cells were mapped on our reference using Symphony ^14^ with a batch correction. Mapped cells were used to assess cell frequency and NMF cell features for each cluster. (B) Bar plots showing the number of samples (left) and the number of CD4^+^ T cells (right) enrolled in the meta-analysis. The dashed line in the left plot indicates a sample size of 10. (C) Dot plot showing changes in cell frequency at cluster L2 resolution. Dot colors show coefficients, and sizes show the significance of the Generalized Linear Model (Methods). Detailed statistics can be found in Table S8. Only significant dots (p_adj_ < 0.05) are shown. (D-F) PCA plots of samples based on cell frequencies. Sample distributions for each disease state (D), loading vectors for each cell type (E), and sample characteristics in healthy donors (F) are shown. (G) Chord diagram showing the top 100 significant associations with positive coefficients between NMF features and cells in each condition, calculated by GLM (Methods). Detailed statistics are shown in Table S9. The thickness of edges indicates the coefficient of GLM, and colors indicate conditions such as diseases, gender, and age. (H) Strategy for predicting autoimmune states from CD4^+^ T cell profiles using machine learning framework. As the input parameters, one model took only cell frequency, age, and gender (without NMFproj), while the other took cell frequency, NMF cell features in Tcm (Th0) and Tnaive, age, and gender (with NMFproj). (I) Receiver operating characteristic (ROC) curves of logistic regression models trained by cell frequencies (top left), by NMFproj values in Tnaive and Tcm (Th0) (top right), and both cell frequencies and NMFproj values (bottom). SLE, COVID-19, and SLE patients were trained on 159, 116, and 35 patients with the same number of healthy subjects, and evaluated on 40, 89, and 17 patients and the same number of healthy subjects from independent data sets. Numbers in parentheses indicate the area under the curve (AUC).

Our meta-analysis could detect these effects as the increase of Tnaive *MX1*. Moreover, in our meta-analysis, Tcm (Th17) was increased in various diseases, including previously reported diseases such as MG ^69^, MS ^70, 71^, and psoriasis ^72^. Regarding Tregs, we reported that Fr. II (CD45RA^-^ CD25^+^) is increased in sarcoidosis, while Fr. I and III are increased in active SLE ^30^. Another group reported an increase of Tregs in pSS ^73^. In concordance with these observations, Treg increased in SLE, neurosarcoidosis, sarcoidosis, and pSS, especially for Treg Eff in neurosarcoidosis and for Treg Act and Treg Eff in SLE patients (Figures 3C, S7B). Interestingly, the acute infection response to COVID-19 showed an increase in Tnaive, whereas influenza infection showed an increase in Temra. We also found age-dependent Tnaive decrease and Temra, Treg Eff increases concordant with previous reports ^74^ (Figure 3C). Sex differences in immunity are critical, especially for autoimmune diseases, because 80% of autoimmune disease occurs in females ^75^. Previous reports have addressed several changes in females, such as the increase of recent thymic emigrants ^76^ and greater activation responses by *in vitro* stimulation ^77^. As for gender differences, we observed a decrease in Tcm (Th2), Tem (Th1/17), Temra (Th1), and Treg Eff and an increase in Tnaive Act, Tnaive *MX1*, Tnaive *SOX4*, Tem (Tph), and Treg Naive in females. These alterations depending on diseases, gender, and age were also observed as specific distributions on the PCA plot (Figures 3D-F, S7C,D). Overall, we profiled numerical features of CD4^+^ T cells in broad autoimmune status, age, and gender.

Next, to measure the quality changes in autoimmune diseases, we applied NMFproj to the datasets and investigated NMF cell feature changes in each cluster L2 population (Figures 3G, S8B, S9; Table S9). The strongest skews were enriched in NMF7 (INF-F) in SLE and COVID-19 patients in a cell-type-wide manner. We also found that even neutral populations such as Tnaive, Tnaive (Act), and Tcm (Th0) showed disease-specific propensities. For example, NMF0 (Cytotoxic-F) increased in RA, MS, and pSS, NMF10 (Tissue-F) was increased in MS, COVID-19, SLE, and neurosarcoidosis, while NMF3 (Naive-F) decreased in a broad range of autoimmune diseases. In Treg cells, NMF1 (Treg-F) decreased in T1D, MG, and MS, indicating the dysfunction of Treg in these diseases independently of the number of Treg cells. Age-dependent increases of NMF8 (Cent. Mem.-F) and NMF4 (Act-F) were also observed. In females, NMF4 (Act-F) was enhanced broadly. The results of qualitative and quantitative changes were consistent with previous reports, demonstrating the robustness of our catalog. For example, it has been reported that among CD4^+^ T cells, an increase of CXCR4^+^ Tnaive is the dominant change in COVID-19 infection ^78^. This is concordant with the result of our meta-analysis showing an increase in Tnaive and an increase in NMF10 (Tissue-F), which contains *CXCR4* as the feature gene (Table S4), specifically in Tnaive cells. Other findings, such as the upregulation of an activation molecule, CD69, which was the feature gene of NMF10 (Tissue-F), in MS and SLE ^79^, and the reduction of effector Tregs and their decreased function in T1D ^80^, are also consistent with our meta-analysis. In conclusion, by meta-analysis, in addition to quantitative changes, we identified qualitative changes depending on the disease, sex, and age at a resolution that is difficult to observe in existing methodologies.

### Autoimmune states are predictable only from peripheral CD4^+^ T profiles

Given that each autoimmune disease disorder possessed a characteristic CD4^+^ T cell profile, we hypothesized that disease status might be predicted solely from CD4^+^ T profiles by utilizing machine learning techniques and our autoimmune-wide scRNA-seq dataset. To confirm this, we created three models for the prediction of disease status. The first model took the frequency of each cell population in cluster L2, the second model took the cellular features of the NMF in subsets, and the last model took both as input parameters (Figure 3H). We note that, for NMFproj features, only Tnaive and Tcm (Th0) cell features, which were affected by various conditions as discussed previously (Figures 3G, S8A), were used to avoid over-fitting. First, binary classification models were constructed for SLE, COVID-19, and MS to distinguish between certain diseases and healthy individuals using logistic regression where equal numbers of diseased and healthy subjects were used for the training, and the models were evaluated using samples from independent projects from these used for the training (Figure 3I). SLE and COVID-19 yielded relatively good predictions from cell frequencies alone (AUC: 0.84, 0.82 in SLE and COVID-19, respectively). NMF cell features alone also could predict SLE and COVID-19 well (AUC: 0.94, 0.85 in SLE and COVID-19, respectively), and the perdition using both cell frequencies and NMFproj values further improved accuracy (AUC: 0.94, 0.91 in SLE and COVID-19 respectively). For MS, for which hematological biomarkers have not yet been well-established, prediction from cell frequencies or from NMFproj values alone was not successful (AUC: 0.61, 0.47 in cell frequency and NMFproj respectively), while both cell frequencies and NMFproj values resulted in better predictions (AUC: 0.75) than previous reports ^81^. Next, we built multiclass classification models that more closely resemble real-world clinical practice. The multiclass classification was assumed to be a more difficult task due to the similarities among autoimmune diseases and the imbalance of training sample sizes. We trained a gradient boosting model with 5-fold cross-validation on data from 714 samples from 8 diseases or healthy for which at least ten samples were available for training and then evaluated the model using the data from the independent dataset (Figures S8B,C). The model trained only by cell frequencies or by NMFproj values could predict COVID-19, SLE, HC, and MS (Area Under the Precision-Recall Curve (PR-AUC): 0.72, 0.68, 0.48, 0.12 in the cell frequency model and 0.91, 0.75, 0.47, 0.35 in the NMFproj model, for COVID-19, SLE, HC, and MS respectively), and the model trained by both cell frequencies and NMFproj values in Tnaive and Tcm (Th0) was marked with superior accuracy (PR-AUC: 0.92, 0.69, 050, 0.33 for COVID-19, SLE, HC, and MS respectively). These results highlight that the disease-specific changes in CD4^+^ T cells, both in terms of quantitative and qualitative alterations, contribute to the prediction of disease status.

### Partitioned heritability of autoimmune diseases on CD4^+^ T cell NMF features

We examined the association between CD4^+^ T characteristics and genetic factors for each disease and trait. Studies using GWAS statistics have reported associations between autoimmune diseases and immune cells ^1, 82^. In particular, stratified linkage disequilibrium score regression (S-LDSC) has revealed the association between CD4^+^ T cells and autoimmune diseases by stratifying the heritability of polygenic autoimmune diseases by genetic features ^83, 84^. Since we captured elaborate CD4^+^ T cell features, we investigated which of these features were associated with each trait using the S-LDSC framework. In addition to the cell-type specific genes of cluster L2, 12-dimensional features extracted by NMF were used as genetic features. The Roadmap Enhancer-Gene Linking (Roadmap) and Activity-By-Contact (ABC) strategies introduced in the sclinker framework ^85^ were used for linking genes and SNPs. Among 180 traits, autoimmune diseases showed significantly high enrichment in NMF features (p=8.51 x 10^-12^), suggesting autoimmunity is closely associated with CD4^+^ T cells (Figure 4A). Cross-sectional disease association revealed that many diseases, such as Inflammatory bowel disease (IBD), RA, and MG, have an enrichment of heritability on NMF1 (Treg-F) (Figure 4B). By focusing on the most accumulated factors for each disease, we found that autoimmune diseases can be divided into several groups.

**Figure 4.**
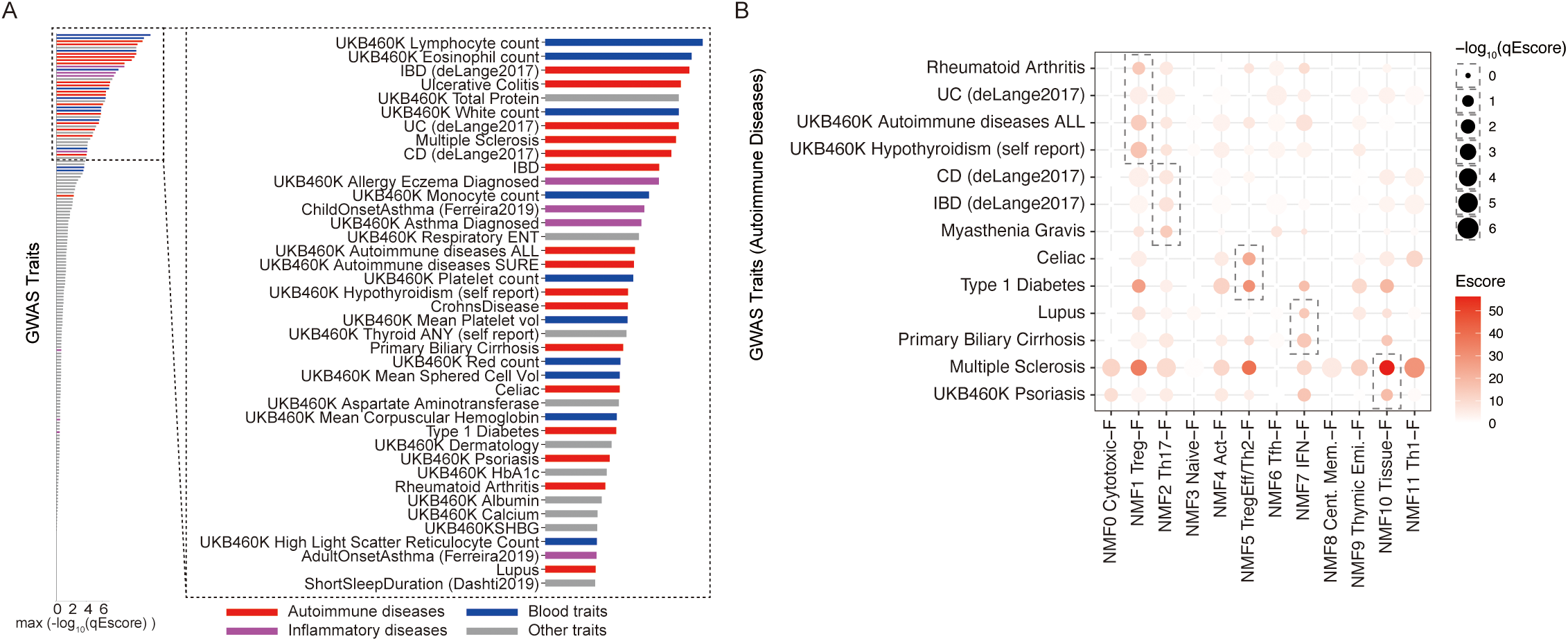
Partitioned heritability of autoimmune diseases by CD4^+^ T cell features. (A) Bar plot showing maximum -log_10_(q_Escore) among NMF gene features. Partitioned heritability was measured using the sclinker framework. Enrichment of each category is the following, Autoimmune diseases: p=8.51 x 10^-12^, inflammatory traits: p=0.131, and blood cell count: p=7.59 x 10^-8^ (Two-sided Mann-Whitney’s U-test). (B) Dot plot showing enrichment of partitioned heritability of autoimmune diseases across NMF gene features. The dashed boxes indicate the factor with the highest Escore for each disease. Duplicated traits were removed for the visualization. Full statistics are shown in Table S11.

For each disease, the most enriched gene features were observed as; NMF1 (Treg-F): RA, UC (deLange), hypothyroidism; NMF2 (Th17-F): CD (deLange), IBD (deLange), MG; NMF5 (TregEff/Th2-F): celiac disease, T1D; NMF7 (IFN-F): SLE, primary biliary cirrhosis; NMF10 (Tissue-F): MS, psoriasis. In MS, accumulation was observed in various features, including NMF2 (Th17-F) and NMF11 (Th1-F). The heritability of each autoimmune disease was accumulated in several factors, suggesting that autoimmune diseases have multiple susceptibilities. In other traits, weak accumulation on NMF1 (Treg-F), NMF2 (Th17-F), and NMF11 (Th1-F) was common in COVID-19 in both severe symptoms and infection, while NMF7 (IFN-F) was infection-specific (Figure S10A). Lymphocyte counts were also susceptible to NMF4 (Act-F) (Figure S10A). We also examined enrichment in cell-type-specific genes. Similar enrichment patterns, such as Treg Naive in most autoimmune diseases and Tnaive *MX1* in SLE, were observed, while caution should be taken as marker gene detection is not optimal for CD4^+^ T cells as in the previous section (Figure S10B). Taken together, we comprehensively profiled heritability enrichment on CD4^+^ T cell gene features across autoimmune diseases.

### Partitioned heritability is associated with qualitative and/or quantitative changes in CD4^+^ T in a disease-specific manner

Lastly, we compared partitioned heritability and observed changes in CD4^+^ T in terms of quantity (cell frequency) and quality (NMFproj) to assess the genetic effect on phenotypic changes (Figure 5A). We first investigated the correlation between enriched heritability and changes in cell frequency and NMF features for diseases that were enrolled in our analysis for both GWAS and meta-analysis. We found several patterns depending on the disease (Figure 5B). First, in MG and psoriasis, both changes in cell frequency and NMF feature correlated with heritability accumulation. In severe COVID-19 (COVID19-A, COVID19-B) and RA, the correlation was mainly observed with changes in quality only. SLE, celiac disease, UC, and SARS-CoV-2 infection (COVID19-C) showed poor correlation with cell frequency.

**Figure 5.**
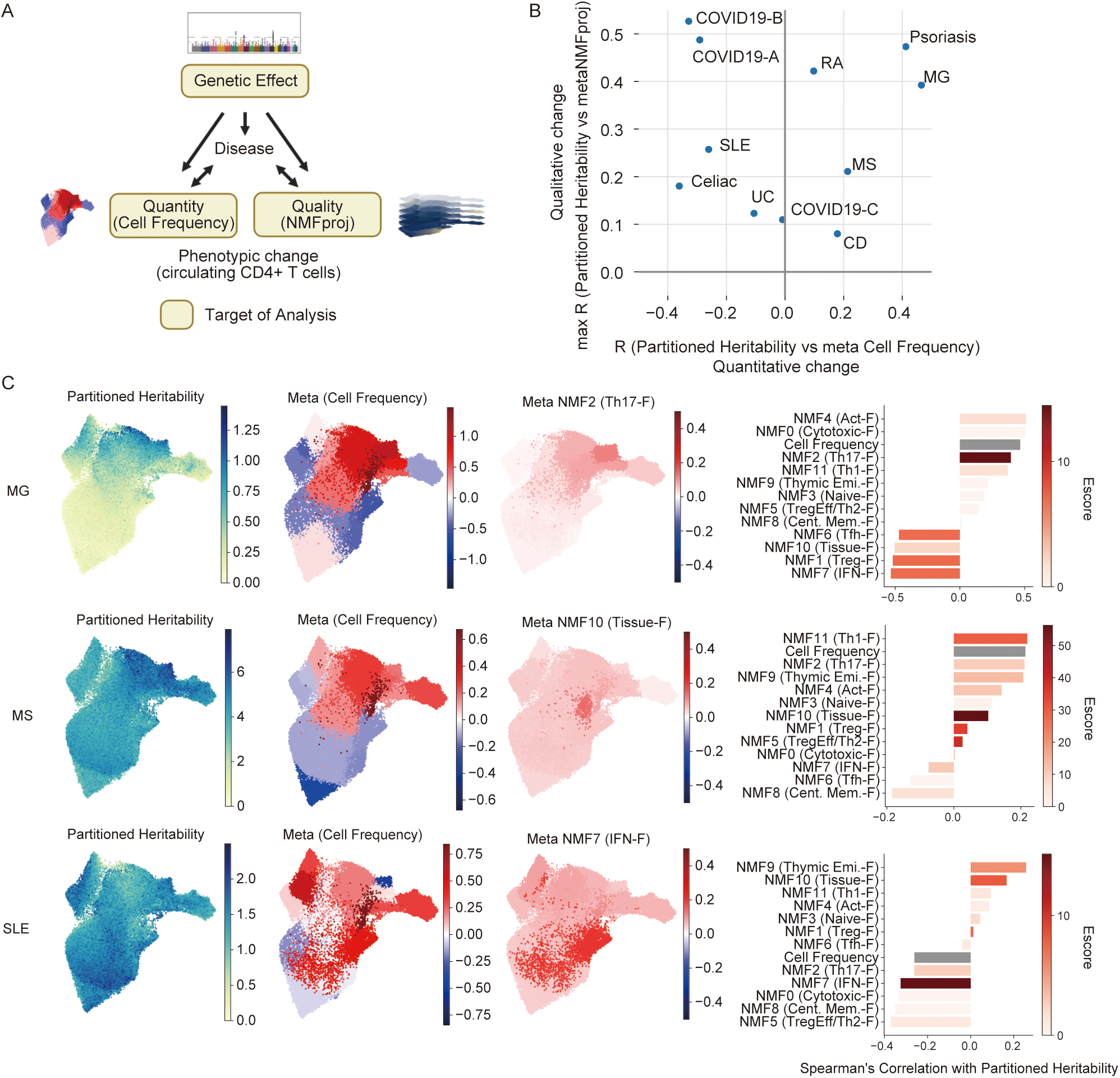
Relationship between genetic factors and phenotypic changes in CD4^+^ T cells. (A) Model of genetic effect on phenotypic changes in CD4^+^ T cells. CD4^+^ T cell changes are observed as qualitative (NMFproj cell features) and quantitative (cell-type frequencies) changes. (B) Scatter plot showing the genetic effect on cell frequencies (x-axis) and NMF features (y-axis). Sclinker weight per cell was calculated by dot products of sclinker outcome (NMF) and NMF cell features. For cell frequencies and NMF cell features, coefficients of GLM output for each cluster L2 population were used. Spearman’s correlation of sclinker weight and cell frequency / NMF cell feature changes were calculated. For the correlation with sclinker and NMF cell feature changes, we used the maximum R among NMF features for the visualization. COVID19-A : Very severe respiratory symptom, COVID19-B : Hospitalized, COVID19-C: SARS-CoV-2 infection. (C) Individual sclinker weights, cell frequency changes (Coef. for each cluster L2), and NMF cell feature changes in the factor with the highest Escore (Coef. for each cluster L2) of MS, MG, and SLE were visualized on the UMAP embeddings (left panel). For the coefficient of the NMF cell feature changes, only one representative factor with the highest Escore for each disease was shown. The bar plot of Spearman’s correlation of cell frequency and NMFproj changes with partitioned heritability is shown in the right panel. The colors of the bar,s except for cell frequency, indicate Escores calculated using sclinker.

Next, we examined MS, MG, and SLE, whose samples were collected in this study, in more detail (Figure 5C). In MG, NMF2 (Th17-F), which accumulated heritability most significantly, was increased in Tcm (Th17) and Tem (Th1/17) in correlation with genetic factor accumulation, and the cell frequency was also correlated with genetic factors (Figures 5C, S9). On the other hand, NMF1 (Treg-F), which showed the next highest accumulation of heritability, was negatively correlated with the genetic effect and lower in Treg cells (Figure S9), indicating that the dysfunction of Treg cells might be enhanced by the genetic effect. In MS, the highest heritability accumulation was observed in NMF10 (Tissue-F). This factor was increased in all cell populations without cell specificity, resulting in a low correlation with the heritability. SLE susceptibility was most accumulated in NMF7 (IFN-F). In our meta-analysis, an enhancement of NMF7 (IFN-F) was observed in all cell populations, especially in Tnaive *MX1*. Overall, our study cataloged heritability enrichment and phenotypic changes across autoimmune diseases, enabling elucidation of the disease-specific effect of underlying genetic factors on CD4^+^ T cell phenotypes.

## Discussion

The classifications and characterizations of CD4^+^ T cells have been challenging, with cellular heterogeneity being a major obstacle ^4^. In this study, by performing single-cell analysis on CD4^+^ T cells from autoimmune and healthy subjects, we succeeded in mutually exclusive and collectively exhaustive subtype identifications of peripheral CD4^+^ T cells. Moreover, in contrast to the conventional dualistic comparisons such as Th1 vs. Th2 and Treg vs. Th17, the NMF-based decomposition revealed that CD4^+^ T cells are formed by a combination of 12 features rather than simple contradistinction. While qualitative profiling by NMF was not suitable for numerical evaluation, it allowed for a more robust assessment of gradual cell populations. Moreover, our results can also be extrapolated for other single-cell and bulk RNA-seq studies by using a label transfer and the projection of NMF features.

These analytical frameworks also allowed us to perform autoimmune-wide single-cell meta-analyses and integration of CD4^+^ T cell features with GWAS. As a result, we comprehensively cataloged CD4^+^ T cell alterations in 20 diseases, providing a valuable resource for a broad range of disease research. The assessment of qualitative changes through NMFproj enabled us to explore biological insights, such as Treg functional abnormalities, that were previously unattainable using cytometry. Furthermore, the decomposition of gene programs using NMF was beneficial not only for T-cell profiling but also for interpreting GWAS results. We found that genetic factors can have both disease-specific and cross-disease impacts on autoimmune conditions. The accumulation of heritability on Tregs across diseases and the disease specificity of other features may be potential clues for future therapeutic development.

By examining genetic factors, CD4^+^ T cell changes, and TCR characteristics in a disease-specific manner, we gained valuable insights into various diseases. For example, MG is caused by autoantibodies against the neuromuscular junction, with germinal center responses involving Tfh cells and B cells within the thymus ^7, 86^. While Th17 function enhancement has been reported in MG ^69, 87^, we also observed a heritability enrichment in NMF2 (Th17-F), suggesting that tissue damage by Th17 cells may contribute to symptom completion and persistence. In MS, we observed an increase in NMF10 (Tissue-F) and heritability enrichment in addition to the previously known Th17 and Th1 increase and functional enhancement ^70, 71, 88^. These results suggested that a strong tissue inflammatory response is involved in MS. These results emphasized that the tissue-specific gene program centered on the AP-1 family may be a novel MS-specific therapeutic target ^89^. Additionally, in MS, while Treg Eff slightly increased, the quality of Treg cells in terms of transcriptome and TCR was low, indicating that the compensation of Tregs from Tconvs is an explanation for Treg dysfunction in MS ^90^. In SLE, our analysis of heritability enrichment and qualitative alterations supported the traditional belief that Type I IFN is central to the disease ^91^. Type I IFN drives differentiation into Tregs and Th1 cells ^92, 93^, and our result suggested that the pleiotropic effect of Type I IFN contributed to the complicated cell frequency changes observed in this study. Furthermore, TCR overlaps between Tcm (Tfh) and Treg Act were observed specifically in SLE, suggesting potential Treg-Tfh plasticity in SLE similar to reported Tfh-Treg plasticity under certain inflammatory conditions ^94, 95^. We identified distinct CD4^+^ T cell responses between COVID-19 and influenza infection, with an increase in Tnaive cells in COVID-19 and Temra cells in flu. This divergence may reflect differences between pre-trained immunity to influenza and initial responses to SARS-CoV-2, as most COVID-19 samples were collected before the vaccine rollout. In addition, our meta-analysis revealed sex and age-related CD4^+^ T cell changes, with new observations such as increased Tnaive *MX1* and Tem (Tph) in females, potentially contributing to gender differences in autoimmune disease incidence. Thus, our study highlights the CD4^+^ T cell features of each disease and condition, providing new insights for consideration.

Additionally, this study created a comprehensive CD4^+^ T cell catalog across various diseases for the first time, providing the opportunity to tackle the challenging task of assessing whether disease prediction is feasible using CD4^+^ T cell profiles. The machine learning model showed that disease status could be predicted only from CD4^+^ T profiles. Although we still could not collect samples abundantly for the model training for clinical applications, this study showed the potential for capturing undiagnosed autoimmune diseases from cellular conditions in the future. The predictability also indicated that changes in CD4^+^ T profiles clearly characterized each disease, emphasizing the importance of fine-tuned treatments for individual diseases.

In our discussion, it is important to address a key limitation of our study, which is that it focused on peripheral blood and did not evaluate tissue alterations, such as barrier tissue-specific programs ^96^. This might be the reason that we could not capture some known vital phenomena for CD4^+^ T cells, such as anergy and exhaustion. On the other hand, the NMF defined in peripheral blood fitted well with tissue CD4^+^ T cells and tumor-infiltrating T cells, suggesting that peripheral blood can be used as a snapshot of complex T cell responses and that these profiles are applicable to tissues.

Collectively, we constructed the frameworks for extracting the CD4^+^ T cell programs, enabling a comprehensive interpretation of CD4^+^ T cells. Moreover, the landscape of disease-specific CD4^+^ T cell alterations and genetic effects provides biological insights for potential precision medicine.

## STAR Methods

### Human samples

The study using human samples was reviewed and approved by the Research Ethics Committee of Osaka University and carried out in accordance with the guidelines and regulations. Human samples were collected under approved Osaka University’s review board protocols: ID 708-10. Written informed consent was obtained from all donors.

### Cell preparation and sequencing of scRNA-seq

From blood collected using heparin-coated tubes, we first collected PBMCs using Ficoll-Paque (Cytiva). PBMCs were washed, blocked with Fc Receptor Binding Inhibitor Polyclonal Antibody, Functional Grade, eBioscience™ (Thermo Fisher Scientific), and stained with FITC-labeled anti-CD3 mAb (dilution: 1/100, UCHT1, BD Bioscience), APC-labeled anti-CD4 mAb (dilution: 1/100, RPA-T4, Thermo Fisher Scientific), PE-labeled anti-CD19 mAb (HIB19, BioLegend), Live/Dead (Thermo Fisher Scientific). Live-CD3^+^CD4^+^CD19^−^ cells were isolated using BD Biosciences FACS Aria II or BD Biosciences FACS Aria III. CD4^+^ T cells and B cells were mixed in equal numbers in some samples.

The sorted cells were loaded to Chromium Next GEM Chip G (10x Genomics) on Chromium Controller (10x Genomics) for barcoding and cDNA synthesis. The library construction was performed using Chromium Next GEM Single Cell 5’ Kit v2 and Chromium Single Cell Human TCR Amplification Kit (10x Genomics) for 5’ according to the manufacturer’s protocol. The libraries were sequenced on NovaSeq6000 (Illumina).

### Preprocess of scRNA-seq data

Sequenced reads were processed using Cell Ranger (v4.0.0) with pre-built reference refdata-gex-GRCh38-2020-A and refdata-cellranger-vdj-GRCh38-alts-ensembl-4.0.0 downloaded at 10x GENOMICS’ website. Quantified expressions were preprocessed and visualized using Scanpy 1.8.1 ^97^ and Python 3.8.0. For CD4^+^ T cell and B cell mixed samples, we extracted only CD4^+^ T cells as following procedures. Briefly, we normalized (sc.pp.normalize_total) gene expression, log-transformed it (sc.pp.log1p), extracted highly variable genes (HVGs) (sc.pp.highly_variable_genes with min_mean=0.0125, max_mean=3, min_disp=0.5), computed PCA (sc.tl.pca) and neighbors (sc.pl.neighbors with n_neighbors=10, n_pcs=40), computed clusters using Leiden algorithm (sc.tl.leiden), and embedded using UMAP algorithm (sc.tl.umap). CD3E-positive and MS4A1-negative clusters were extracted as CD4^+^ T cells and used for the analysis. Cells with mitochondrial genes were higher than 10%, detected genes less than 200, or annotated as multichain by scirpy were filtered out. Variable genes of TCR alpha and beta were removed for the clustering and embedding to remove the effect of clonal expansion. Gene expressions were preprocessed by sc.pp.normalize_per_cell with counts_per_cell_after=1e4, sc.pp.log1pp, retained HVGs. The inference of the cell cycle was performed using the sc.tl.score_genes_cell_cycle function following the tutorial (https://nbviewer.jupyter.org/github/theislab/scanpy_usage/blob/master/180209_cell_cycle/cell_cycle.ipynb). Total counts of UMI, % mitochondrial genes, S score, G2M score were regressed out using sc.tl.regress_out and scaled using sc.tl.scale. Then, principal components were computed using sc.tl.pca. The batch effect of samples was removed by the Harmony algorithm ^13^. Cells were embedded by UMAP using sc.tl.umap (spread=1.5), and clustered using sc.tl.leiden (resolution=1.2). Re-clustering and embedding were performed using sc.tl.umap (spread=1.5), clustered using sc.tl.leiden (resolution=1.7) after removing clusters containing doublets with B cells, monocyte lineages, etc. We defined cluster L1 as a large classification using the leiden clusters. Next, for some clusters, concatenation or re-clustering was performed with sc.tl.leiden (resolution 0.3-1) to divide clusters at the minimum resolution with distinct marker genes. We defined cluster L2 as a smaller classification. Marker genes were determined using sc.tl.rank_genes_groups with method=’t- test_overestim_var’.

### Integration with bulk RNA-seq dataset

Fastq files were processed using an RNA-seq integrative pipeline, ikra (v2.0.1) ^98^, composed of Trim Galore! 0.6.7 ^99^, Salmon 1.4.0 ^100^, tximport 1.6.0 ^101^ with the reference Gencode M26 for mice and 37 for humans. Datasets for which the TPM matrix was provided were downloaded and used directly for analyses. For the ImmuNexUT (E-GEAD-397) dataset, the downloaded count matrix was converted to TPM.

For the correlations between bulk RNA-seq and scRNA-seq datasets, TPM or scaledTPM expression matrix of bulk RNA-seq were normalized using sc.pp.normalize_per_cell (counts_per_cell_after=1e4), sc.pp.log1p, sc.pp.scale (max_value=10), concatenated to scRNA-seq object, and calculated the correlations using sc.tl.dendrogram with the default parameters.

### Gene expression decomposition using NMF

To decompose cellular processes, we applied NMF implemented in scikit-learn (v0.24.2) to normalize gene expression of HVGs. Using NMF, the normalized gene expression X was decomposed as follows;

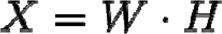

where W and H possess n components. In the analysis, we set the number of components as 12 based on the two criteria; i) more than the elbow in the distribution of explained variance, ii) before the jump up of the maximum inter-components Spearman’s correlation. The explained variance was calculated as follows;

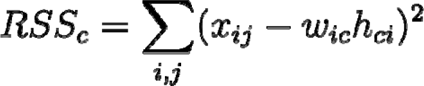

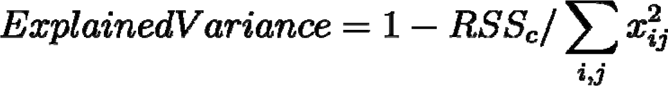

For the pathway enrichment analysis, we extracted the top 100 genes with the highest feature value for each component and converted gene symbols into Entrezid using the bitr function provided by clusterProfiler (3.16.1) ^102^ and computed enriched Reactome pathways using the compareCluster function of clusterProfiler with the enrichPathway function in ReactomePA (1.32.0). For the projection of gene features defined by NMF, we performed NMF with pre-computed W using scikit-learn. The matrix W was converted to mouse genes using a list of human and mouse homologs provided at http://www.informatics.jax.org/homology.shtml. For genes with multiple homologs, one of the genes was retained. For the NMF calculation, only overlapped genes were used. To examine whether the selected HVGs of fixed W can capture HVGs in a query dataset, we calculated the proportion of the number of HVGs included in fixed W against the number of HVGs in the query dataset as POH. In the CD4^+^ T data set, we determined that if the POH is below 0.1, which is the conservative threshold from the distribution of the null hypothesis (Figure S3C), there is a variance that cannot be represented by the NMFproj. sc.pp.highly_variable_genes in scanpy with the following parameters; min_mean=0.0125, max_mean=3, min_disp=0.1 was used for the calculation of HVGs of the query datasets and selected top 500 genes regarding normalized dispersion ^103^ with the exclusion of VDJ genes of TCR and IG. A framework for NMF projection is available at https://github.com/yyoshiaki/NMFprojection.

For the analysis of ImmuNexUT, the count data was downloaded at (https://humandbs.biosciencedbc.jp/en/hum0214-v3) and converted to TPM. We removed BD, AAV, AOSD, and SjS samples from the analysis because these samples are processed in a different procedure from other samples. TPM matrix was decomposed with the pre-computed gene feature matrix using NMFproj. Extracted NMF feature H was tested using a multiple linear regression provided as the formula.api.ols function by a python package statsmodels (0.12.0) with a model, NMF_i ∼ Disease + Age + Gender + 1.

### TCR analysis

For TCR analysis, we used the standard pipeline of scirpy 0.10.1 ^104^ according to the official tutorial (https://scverse.org/scirpy/latest/tutorials/tutorial_3k_tcr.html). Briefly, clones were defined by clonotypes using scirpy.tl.define_clonotypes with parameters; receptor_arms=“all”, dual_ir=“primary_only”. Repertoire similarities were measured using the function scirpy.tl.repatoire_overlap. TiRP score was calculated according to the instruction in the repository (https://github.com/immunogenomics/TiRP.git).

### Heritability partitioning

To assess the contribution of each cell-type-specific gene expression and the NMF component, we applied S-LDSC with Roadmap ABC-immune enhancer-gene linking strategy implemented using an sc-linker pipeline ^85^ with slight modifications (https://github.com/yyoshiaki/sclinker-skg). We only used HVGs defined by the preprocessing section in the analysis. We used min-max scaled gene scores for the cell-type gene programs as the gene weights. For NMF components, we used the gene feature matrix W with the min-max scaling for the gene weights. Using the gene weights, LD scores for each category were calculated with European LD scores used in the article ^85^ and Roadmap_U_ABC for blood (https://storage.googleapis.com/broad-alkesgroup-public/LDSCORE/Dey_Enhancer_MasterReg/processed_data, https://storage.googleapis.com/broad-alkesgroup-public/LDSCORE/DeepLearning/Dey_DeepBoost_Imperio/data_extra/AllPredictions.AvgHiC.ABC0.015.minus150.withcolnames.ForABCPaper.txt.gz). In addition to the sumstats files provided in gs://broad-alkesgroup-public/LDSCORE/all_sumstats, we used several additional sumstats by processing using munge_sumstats.py in LDSC v1.0.1 (Table S10). S-LDSC was performed with the baseline-LD model v2.1 (https://storage.googleapis.com/broad-alkesgroup-public/LDSCORE/1000G_Phase3_baselineLD_v2.1_ldscores.tgz). The Enrichment score (Escore) was calculated as the difference between the enrichment for annotation in a particular program against an SNP annotation for all protein-coding genes with a predicted enhancer-gene link in the blood. We also used FDR calculated from the p-value of Enrichment outputted by S-LDSC.

### Meta-analysis of CD4^+^ T cells from public datasets

We collected scRNA-seq data from PBMC generated by 10x platforms, Seq-Well or SPLiT-seq (Parse Biosciences WT Mega) ^6, 8, 36–57^ (Table S6). If the count matrix was available, we used the quantified matrix. Otherwise, we quantified the expression using Cell Ranger with pre-built reference refdata-gex-GRCh38-2020-A. As the sample QC, samples with XIST mean expression (count) > 0.05 were inferred as female. If the inferred gender and metadata differed, we removed the sample from the analysis. We extracted CD4^+^ T cells from published data of PBMCs using Azimuth 0.4.4 ^58^. We created Seurat Object using the CreateSeuratObject function implemented in Seurat 4.1.0 ^58^ with parameters min.cells=3, min.features=200. We also filtered out the cells that express≧10% mitochondrial genes in their total gene expression. We normalized the expression using SCTransform with parameters method= “glmGamPoi”, ncells=2000, n_genes=2000, do.correct.umi=FALSE. In this procedure, we used Azimuth reference data v1.0.0 human_pbmc loaded from the website (https://seurat.nygenome.org/azimuth/references/v1.0.0/human_pbmc). We found anchors between query data and Azimuth reference data (FindTransferAnchors with parameters k.filter=NA, normalization.method= “SCT”, dims=1:50, n.trees=20, mapping.score.k=100), transferred cell type labels (TransferData with parameters dims=1:50, n.trees=20) and calculated the embeddings on the reference supervised PCA (IntegrateEmbeddings with the default options) and neighbors (FindNeighbors with parameter l2.norm=TRUE). We transformed an NN index (NNTransform with the default parameters) and projected the query data to the reference UMAP (RunUMAP with the default parameters). We visualized query data by DimPlot, DotPlot, and FeaturePlot.

Next, we mapped extracted cells on our reference using symphony 0.1.0 ^14^ following the vignettes (https://github.com/immunogenomics/symphony/blob/main/vignettes/Seurat.ipynb). First, we created a symphony reference using our dataset. Our scanpy object saved as an h5ad file was converted to h5Seurat using SeuratDisk and loaded as a Seurat object. We used only HVGs for symphony reference to reduce batch effect strictly. Then, the object was preprocessed as follows; SCTransform (method= “glmGamPoi”), ScaleData, RunPCA, RunHarmony.Seurat (group.by= “sample”), FindNeighbors (dims=1:30), RunUMAP2, and buildReferenceFromSeurat. For query mapping, extracted CD4^+^ T cells were normalized (SCTransform with parameter method= “glmGamPoi”) and mapped (mapQuery with parameter do_normalize=FALSE, vars = “batch”) with batch correction against each sample. The cluster L1 and L2 assignments were performed using the knnPredict.Seurat function. We visualized the mapping results by DimPlot and FeaturePlot as the quality control. We are providing the label transfer pipeline at https://github.com/yyoshiaki/screfmapping.

Binominal regression was performed with the formula; (n_cat, n_total - n_cat) ∼ Disease + Age + Gender + Project using the R glm function. For the analysis of enriched NMF components, we first calculated cell profiles using NMFproj with raw counts, and linear regression was performed with the formula; NMF_i ∼ Disease + Age + Gender + Project using the R glm function. For the PCA plot of individuals, we corrected the batch effect by a project by regressing out using GLM with the formula; cell frequency ∼ Disease + Age + Gender + Project. The Chord diagrams were created using pycirclize (0.1.3).

### Machine learning for the prediction of autoimmune states

First, the training and test datasets were split without the study overlap. Cell frequencies and/or NMFproj values in Tcm (Th0) and Tnaive were scaled using StandardScaler (scikit-learn 1.0.2). Note that Tnaive and Tcm(Th0), which have a high degree of nodes in the network, were selected for the NMFproj results to keep the number of parameters low. NMF values were imputed using SimpleImputer (scikit-learn 1.0.2) with parameter strategy=’most_frequent’ trained by training datasets. For the binary classification, we used the LogisticRegression in scikit-learn with the default parameters. For the multiclass classification, the label imbalance was corrected using SMOTE (imbalanced-learn 0.9.1) with parameter sampling_strategy=’all’. Then, LightGBM 3.3.2 ^105^ was used for the model with parameters, ’objective’=’multiclass’ and ’early_stopping_rounds’=10.

### Statical analyses

All statistical analyses were performed in R (4.0.3 or 4.1.2) and Python (3.8.0). FDR was obtained by the Benjamini-Hochberg procedure implemented by a Python package statsmodels (0.12.0). All other statistical analyses are detailed in the respective sections of the article.

## Acknowledgments

This work was supported by Grant-in-Aid for Specially Promoted Research Grant 16H06295 from the Ministry of Education, Culture, Sports, Science, and Technology of Japan and Leading Advanced Projects for Medical Innovation (LEAP, no. 18 gm0010005h0001) from Japan’s Agency for Medical Research and Development (AMED) to S.S. Y.Y. was supported by Takeda Science Foundation. D.T. was supported by the Osaka University Medical Doctor Scientist Training Program. We acknowledge the NGS core facility of the Genome Information Research Center at the Research Institute for Microbial Diseases of Osaka University for its support in RNA sequencing. This work was partly achieved through the use of SQUID at the Cybermedia Center, Osaka University. Some illustrations were generated with BioRender.com. The authors wish to acknowledge Dr. Hirotaka Matsumoto, Nagasaki University for his constructive feedback on NMFproj, Dr. Kazuyoshi Ishigaki, RIKEN for technical advice for TiRP analysis, Nicole Carter and Sarah Schroeder, Parse Biosciences, Dr. Francesca Capon and Dr. Daniel McCluskey, King’s College London, Dr. Karine Chemin, Karolinska institutet, Dr. Lennart M. Roesner, Hannover Medical School (MHH) for sharing single-cell RNA-seq data, and Dr. James Wing for critical comments on the manuscript.

## Data and materials availability

Data sets will be available upon publication. NMFproj is provided at https://github.com/yyoshiaki/NMFprojection. Detailed results of the meta-analysis and sclinker were deposited at https://yyoshiaki.github.io/autoimmune_scRNAseq/Tcells.html.

## Author contributions

Y.Y., D.T., R.M., N.O., and S.S. designed all experiments; Y.N. performed experiments under the supervision of N.O; Y.Y., D.T., and R.M. performed bioinformatics analysis; Y.Y., T.O., M.K., T.M., Y.K., A.K., and H.M. collected samples for analysis; M. W. and X. Z. provided single-cell RNA-seq data; D.M. and D.O. performed library construction and sequencing; Y.T., N.M., and M.A. provided expert advice; Y.Y. prepared the figures; Y.Y., D.T., and R.M. drafted the manuscript; N.O. and S.S. supervised the study; All authors critically reviewed and edited the final version of the manuscript.

## Declaration of interests

The authors declare no competing interests.

## Supplementary Figure Legends

**Figure S1.**
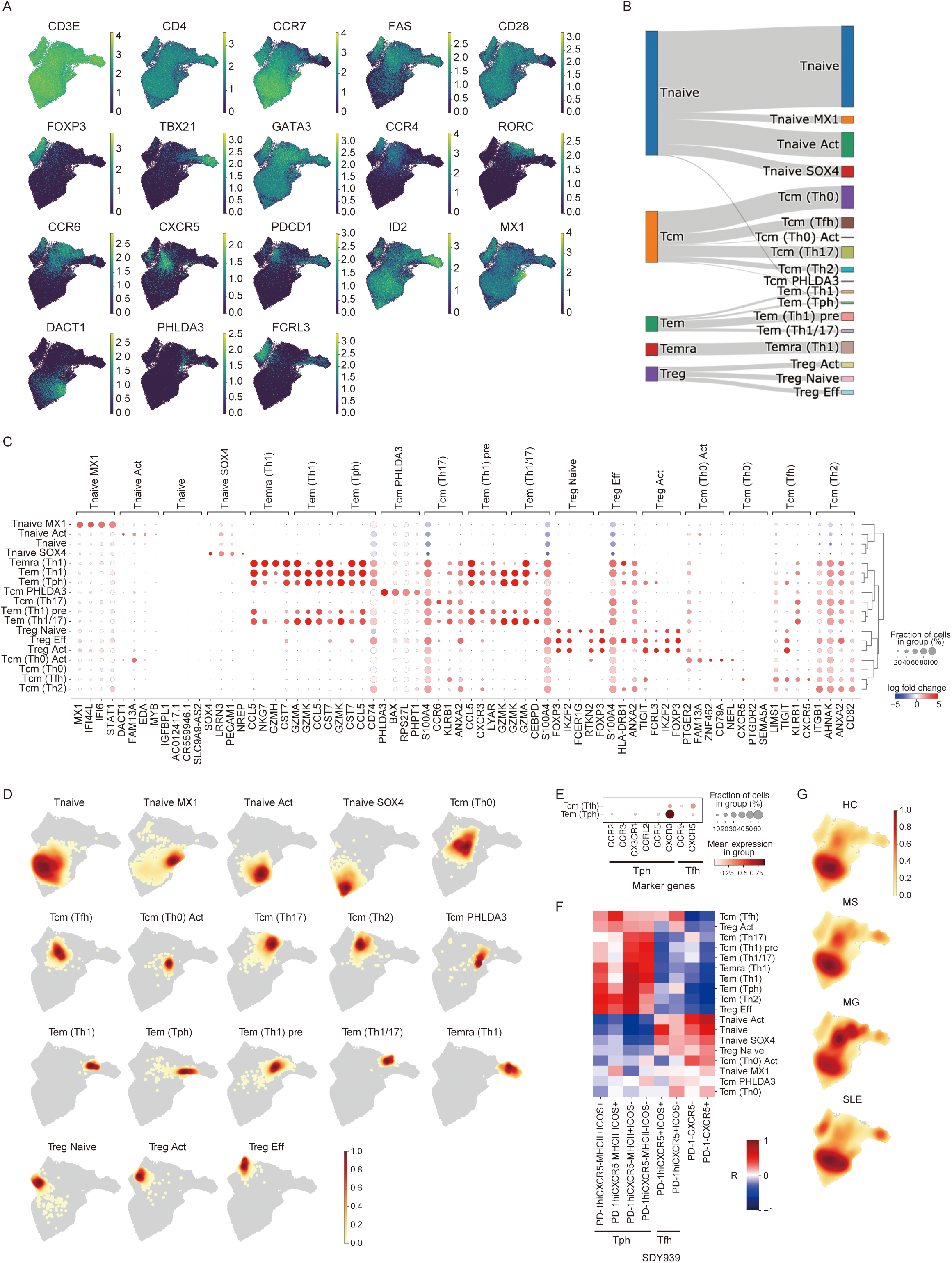
Global characterization of CD4^+^ T cells. (A) UMAP plots depicting gene expressions of marker genes. (B) Sankey diagrams showing cluster assignment of cells in clusters L1 and L2. (C) Dot plot depicting signature genes’ mean expression levels and percentage of cells expressing them across clusters. Marker genes for the plot were calculated by pairwise comparison with a group and the other groups iteratively using scanpy.tl.rank_genes_groups function. (D) Density plot of cell distributions for cluster L2 populations. (E) Dot plot depicting Tph and Tfh marker genes’ mean expression levels and percentage of cells expressing them in Tcm (Tfh) and Tem (Tph). (F) Pearson’s correlation of transcriptome profiles between sorted T cell fractions, including Tph (SDY939) and our scRNA-seq (cluster L2). (G) Density plot of cell distributions for each disease.

**Figure S2.**
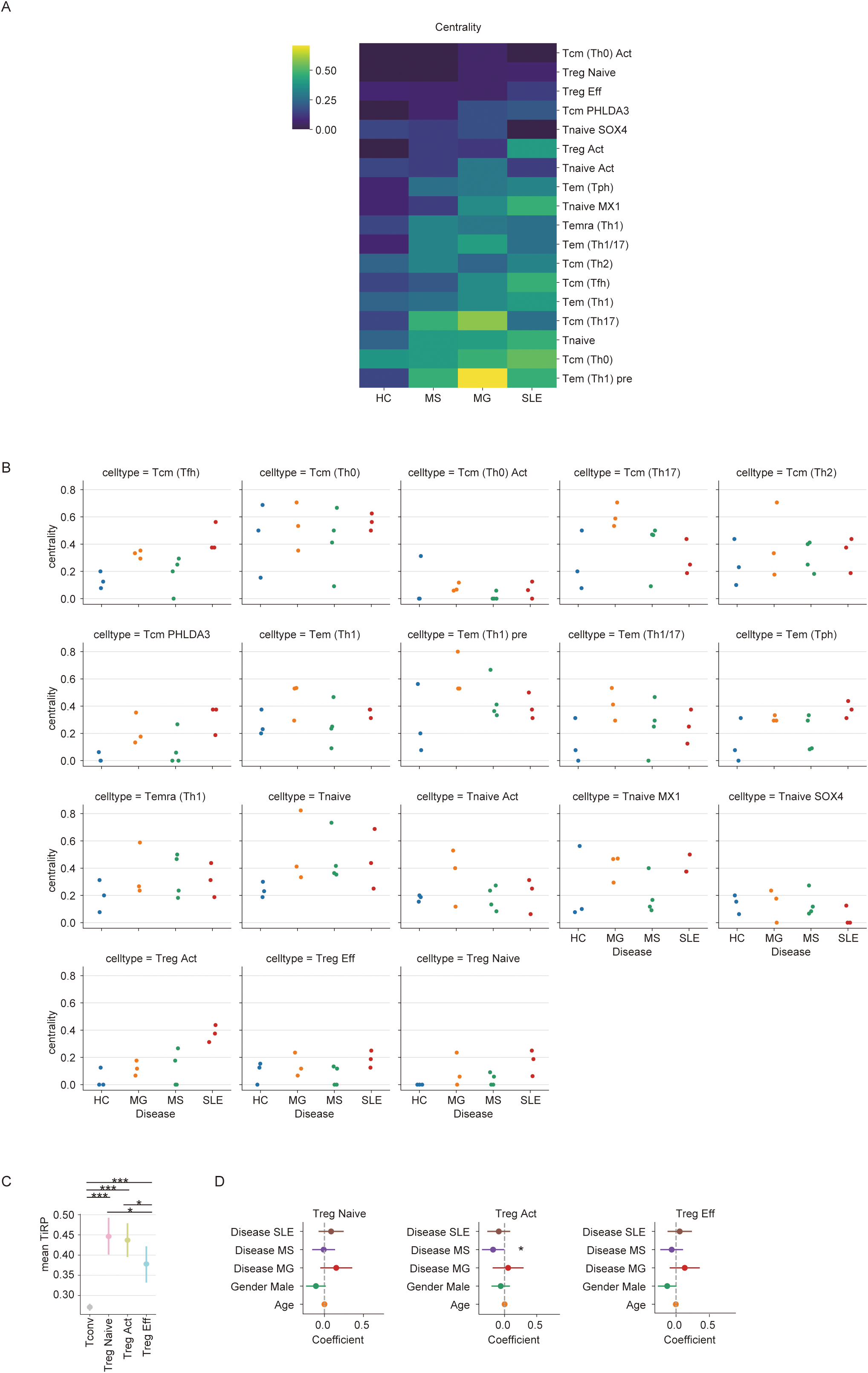
Centralities of TCR networks vary depending on the diseases. (A) Degree centrality of TCR networks for cluster L2. The average of each disease was shown. (B) Individual value of degree centrality of TCR networks. (C) Distribution of mean TiRP scores across Treg clusters. Pairwise Tukey-HSD posthoc tests. The multiple test correction was performed using a two-stage FDR strategy. *: p_adj_ < 0.05, **: p_adj_ < 0.01, ***: p_adj_ < 0.001. (D) Changes in TiRP scores in Treg clusters associated with disease states, age, and sex. The estimated coefficients and the 95 percentiles by multiple linear regression were plotted.

**Figure S3.**
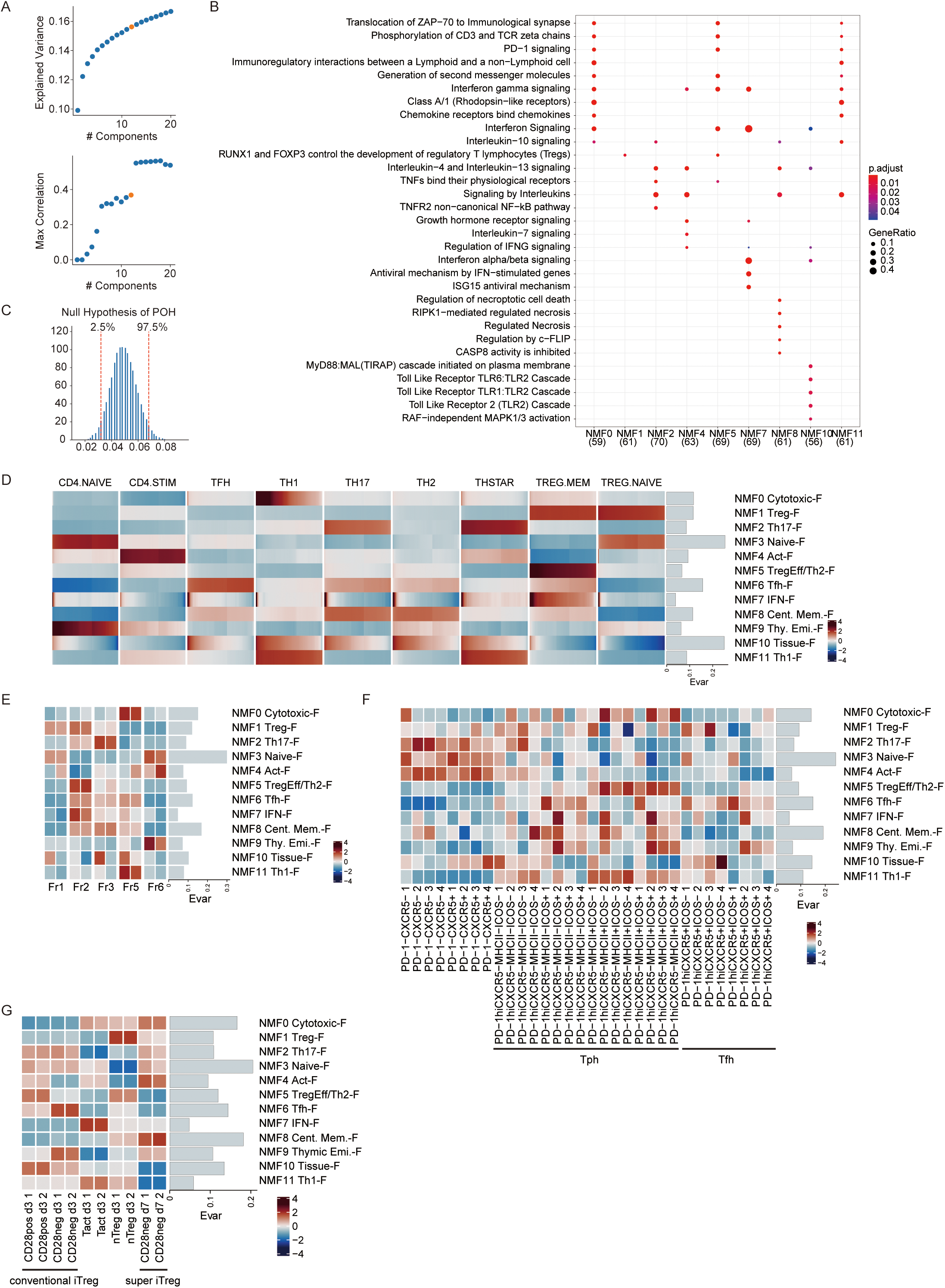
NMF and NMF projection. (A) The statistics for the determination of the number of components. The Y-axis shows explained variance (upper) and maximum correlation of the inter-component (lower). The X-axis shows the number of components. Spearman’s correlation between components of gene features was calculated. (B) Reactome pathways enriched in each gene feature. The dot size indicates the gene ratio or the fraction of genes found in the gene set, and the color indicates p_adj_. (C) Histogram of POH under the null hypothesis for this study setting. We randomly sampled 5000 POH in the null hypothesis calculated from the overlap between random 500 (number of HVGs for the calculation of POH) genes and 1271 (number of HVGs of CD4^+^ T cell) genes. Red dashed lines show 2.5 and 97.5 percentiles. (D) Heatmap showing NMF values of DICE bulk RNA-seq datasets of sorted CD4^+^ T cell fractions. Explained variance (Evar) was also shown on the right side. (E) Heatmap showing NMF values of sorted CD4^+^ T cell fractions by Miyara’s classification (JGAD000214). Explained variance (Evar) was also shown on the right side. (F) Heatmap showing NMF values of sorted Tph fractions (SDY939). Explained variance (Evar) was also shown on the right side. (G) Heatmap showing NMF values of iTreg cells cultured in different conditions (DRA008294). Explained variance (Evar) was also shown on the right side.

**Figure S4.**
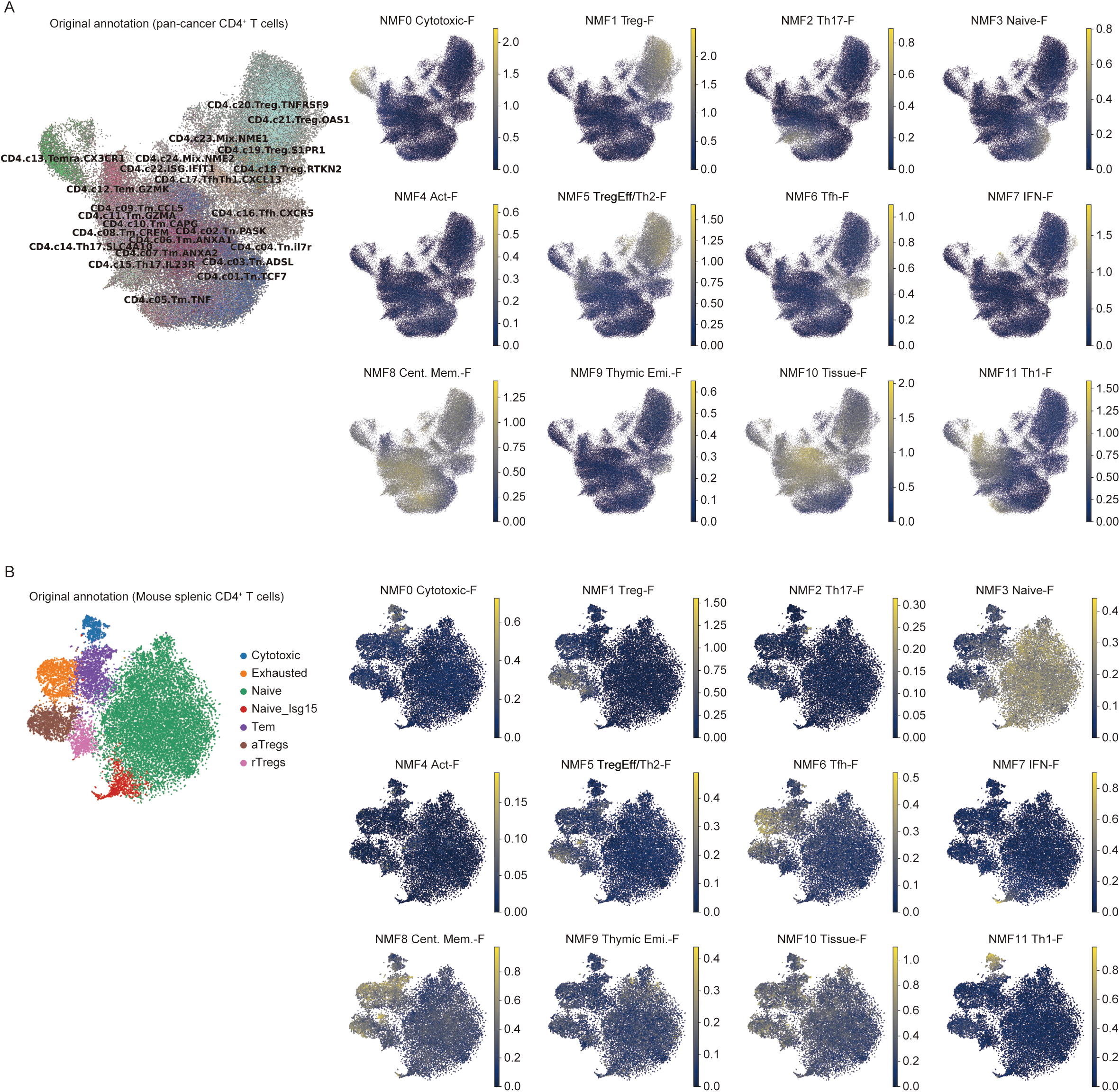
NMFproj applications in tumor-infiltrating T cells and mouse splenocytes. (A and B) UMAP plots showing original cell types (left) and projected NMF cell feature values (right) in pan-cancer tumor-infiltrating T cells scRNA-seq data (GSE156728) (A) and mouse splenic CD4^+^ T cells (SCP490) (B).

**Figure S5.**
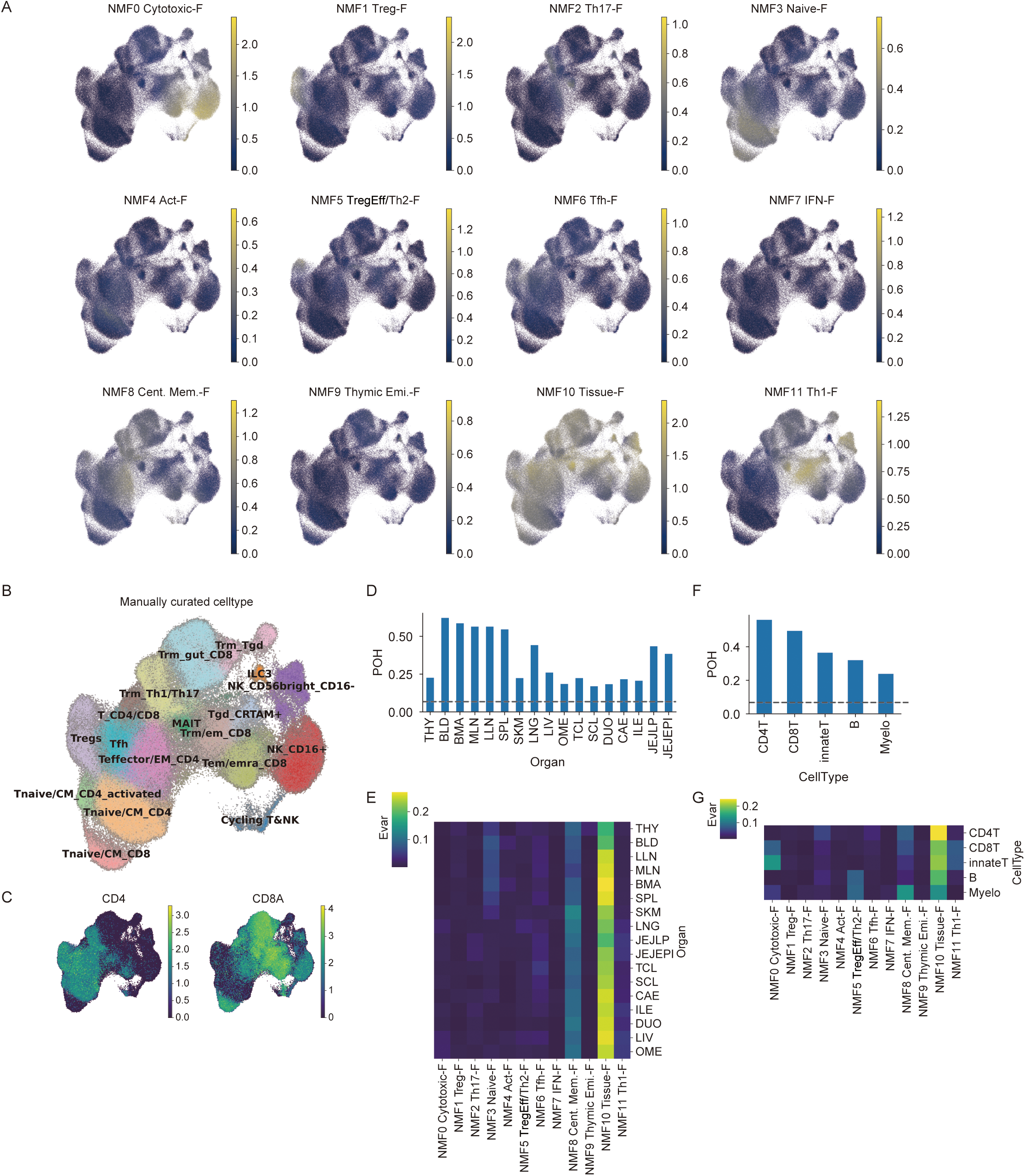
NMFproj contributes to interpreting cross-tissue T cells. (A) Projected NMF cell feature value of cross-tissue T cells scRNA-seq datasets on the UMAP plots. The T & innate lymphoid cells dataset was used for the analysis (https://www.tissueimmunecellatlas.org/). (B and C) Original cell types (B) and the expression of CD4 and CD8A (C) were shown on the UMAP plots. (D and E) Distribution of POH (D) and Evar (E) in each tissue. THY: Thymus, BLD: Blood, BMA: Bone marrow, MLN: Mesenchymal lymph nodes, LLN: Lung-draining lymph nodes, SPL: Spleen, SKM: Skeletal muscle, LNG: Lung, LIV: Liver, OME: Omentum, TCL: Transverse colon, SCL: Sigmoid colon, DUO: Duodenum, CAE: Caecum, ILE: Ileum, JEJLP: Jejunum lamina propria, JEJEPI: Jejunum epithelial. The dashed line indicates the 97.5 percentile of simulated null distribution (Fig. S3C).\ (F and G) Distribution of POH (F) and Evar (G) in each cell type. B and Myeloid cells were also added to the analysis. The dashed line indicates the 97.5 percentile of simulated null distribution (Fig. S3C).

**Figure S6.**
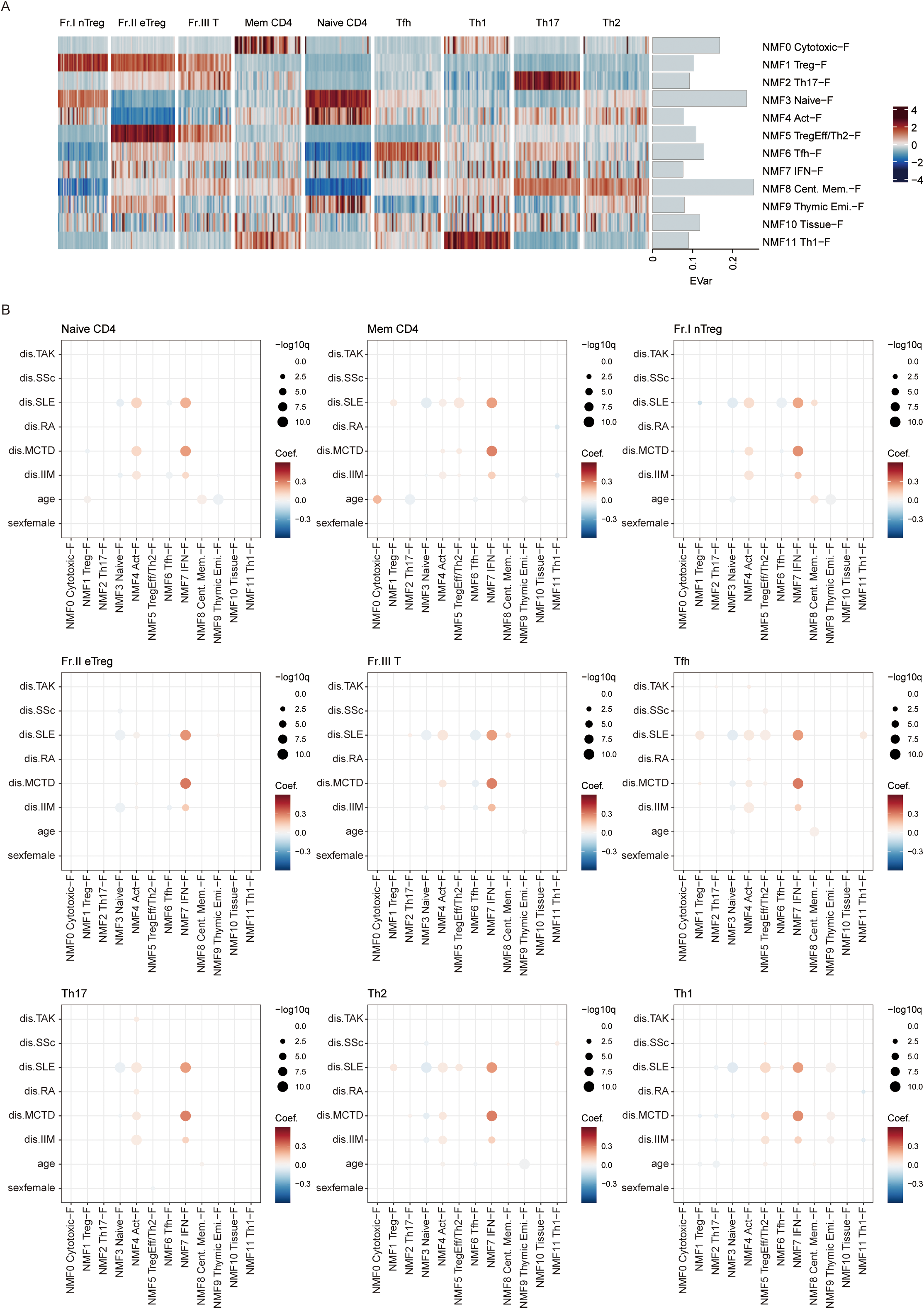
NMFproj reveals disease-specific qualitative changes. (A) Heatmap showing NMF values of sorted CD4^+^ T cell fractions collected from autoimmune patients (E-GEAD-397). Explained variance (Evar) was also shown on the right side. (B) Dot plot depicting NMF cell feature changes in each cell type in E-GEAD-397. Dot colors show coefficients, and sizes show the significance of GLM. GLM was performed with a model, cell frequency, or NMF cell feature ∼ disease + age + gender. IIM: idiopathic inflammatory myopathy, MCTD: mixed connective tissue disease, RA: rheumatoid arthritis, SLE: systemic lupus erythematosus, SSc: systemic sclerosis, TAK: Takayasu arteritis.

**Figure S7.**
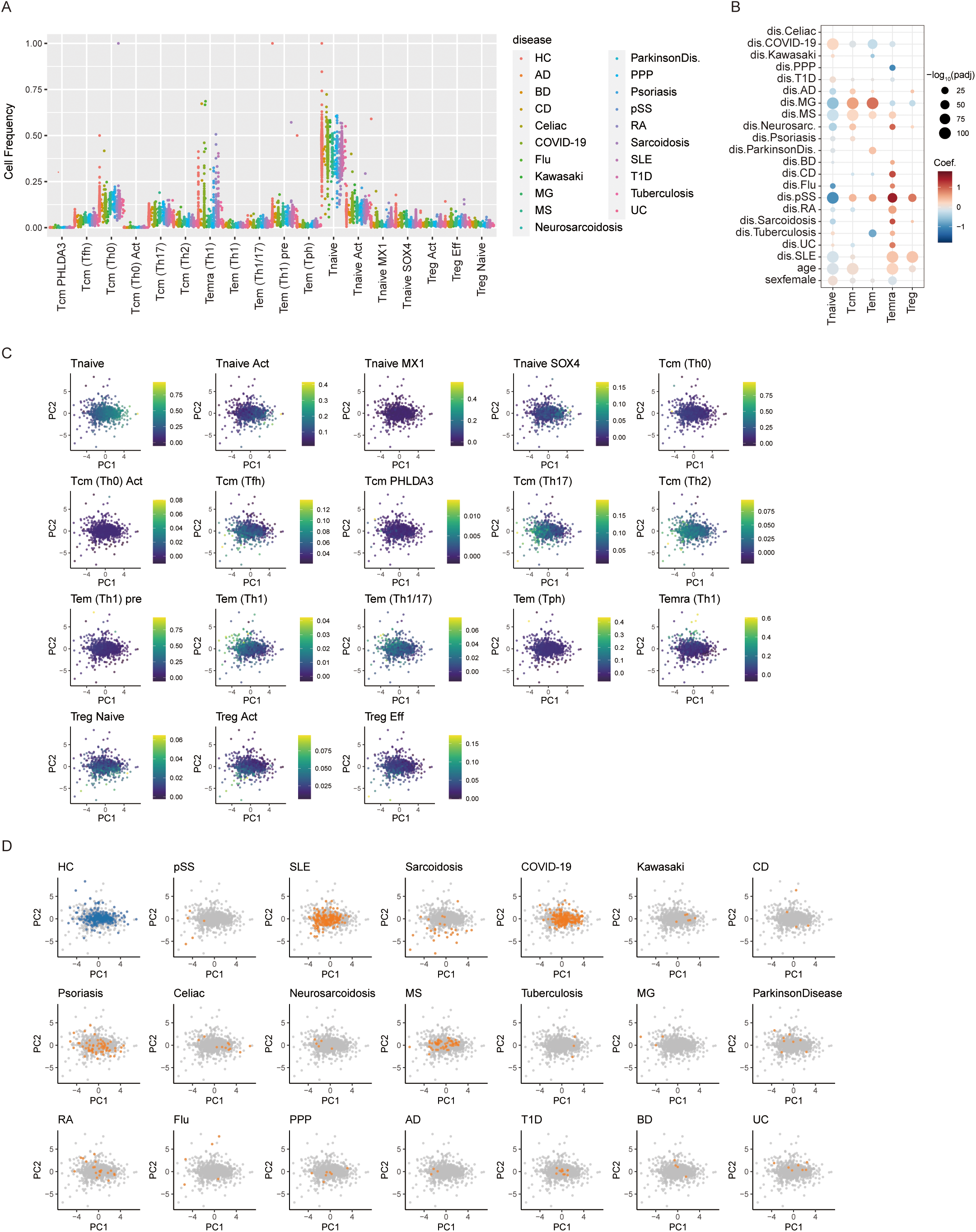
Quantitative alterations revealed by meta-analysis. (A) Swarm plot showing frequencies of cell types in each sample. (B) Dot plot showing changes in cell frequency at cluster L1 resolution. Dot colors show coefficients, and sizes show the significance of the Generalized Linear Model (Methods). Detailed statistics can be found in Table S7. Only significant dots (p_adj_ < 0.05) are shown. (C) Cell frequencies of each population are shown on the PCA plots. (D) Distribution of samples for each disease.

**Figure S8.**
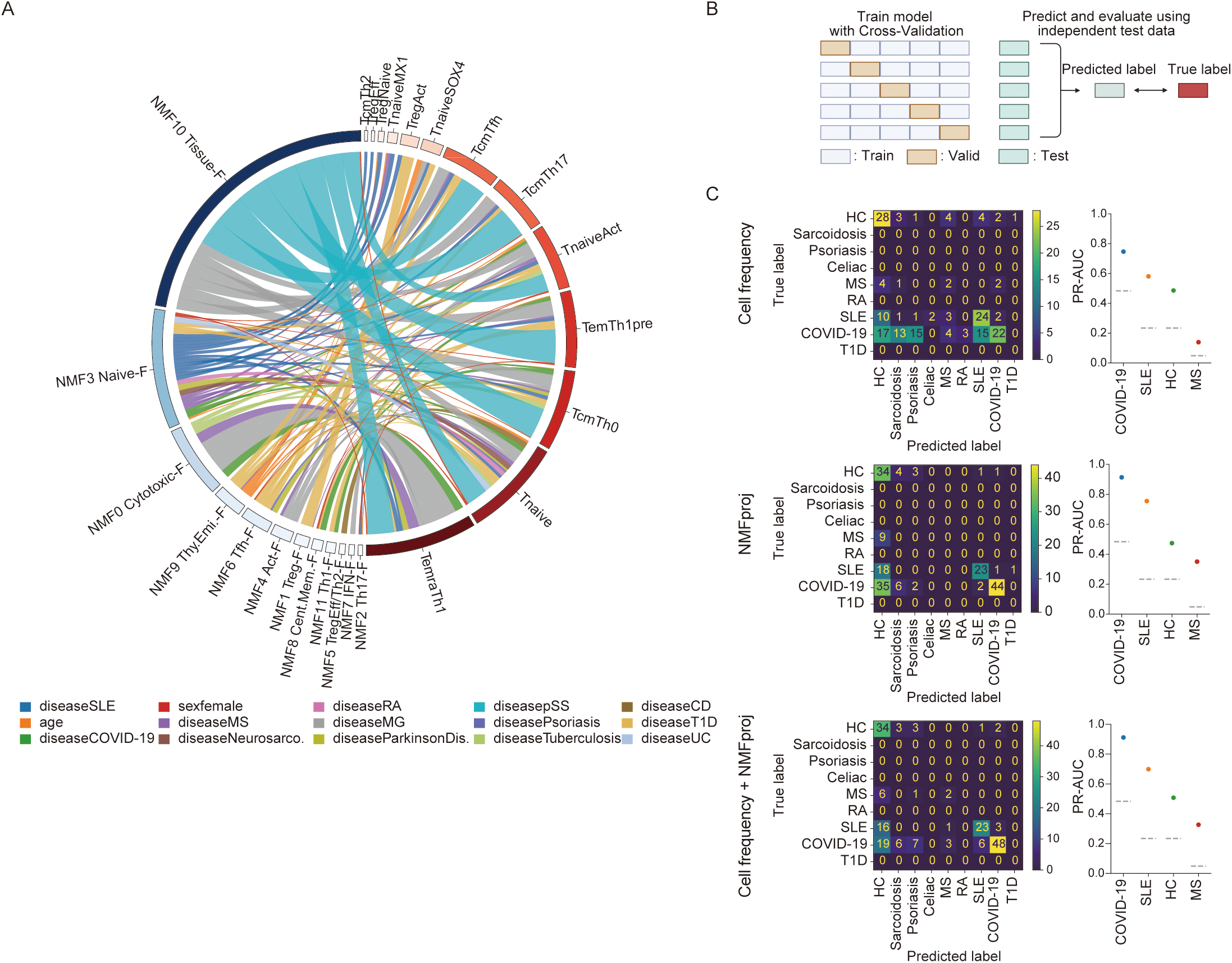
Alterations in CD4^+^ T cells revealed by meta-analysis. (A) Chord diagram showing the top 100 significant associations with negative coefficients between NMF features and cells in each condition, calculated by GLM (Methods). Detailed statistics are shown in Table S9. The thickness of edges indicates the absolute value of the coefficient of GLM, and colors indicate conditions such as diseases, gender, and age. (B) Strategy for multiclass classification by machine learning. The training was performed with cross-validation. The evaluation was performed using the independent dataset of training datasets. (C) Evaluations of models trained by cell frequencies (upper panel), by NMFproj values in Tnaive and Tcm (Th0) (middle panel), and by both cell frequencies and NMFproj values (lower panel). The confusion matrix (left) and PR-AUC (right) are shown. The dashed lines in the PR-AUC plot show the expected PR-AUC scores in random models. The number of samples used for the training is 263, 27, 62, 11, 43, 20, 156, 116, 11 subjects for HC, sarcoidosis, psoriasis, celiac disease, MS, RA, SLE, COVID-19, and T1D, and evaluated on 89, 43, 43, and 9 subjects from independent data sets.

**Figure S9.**
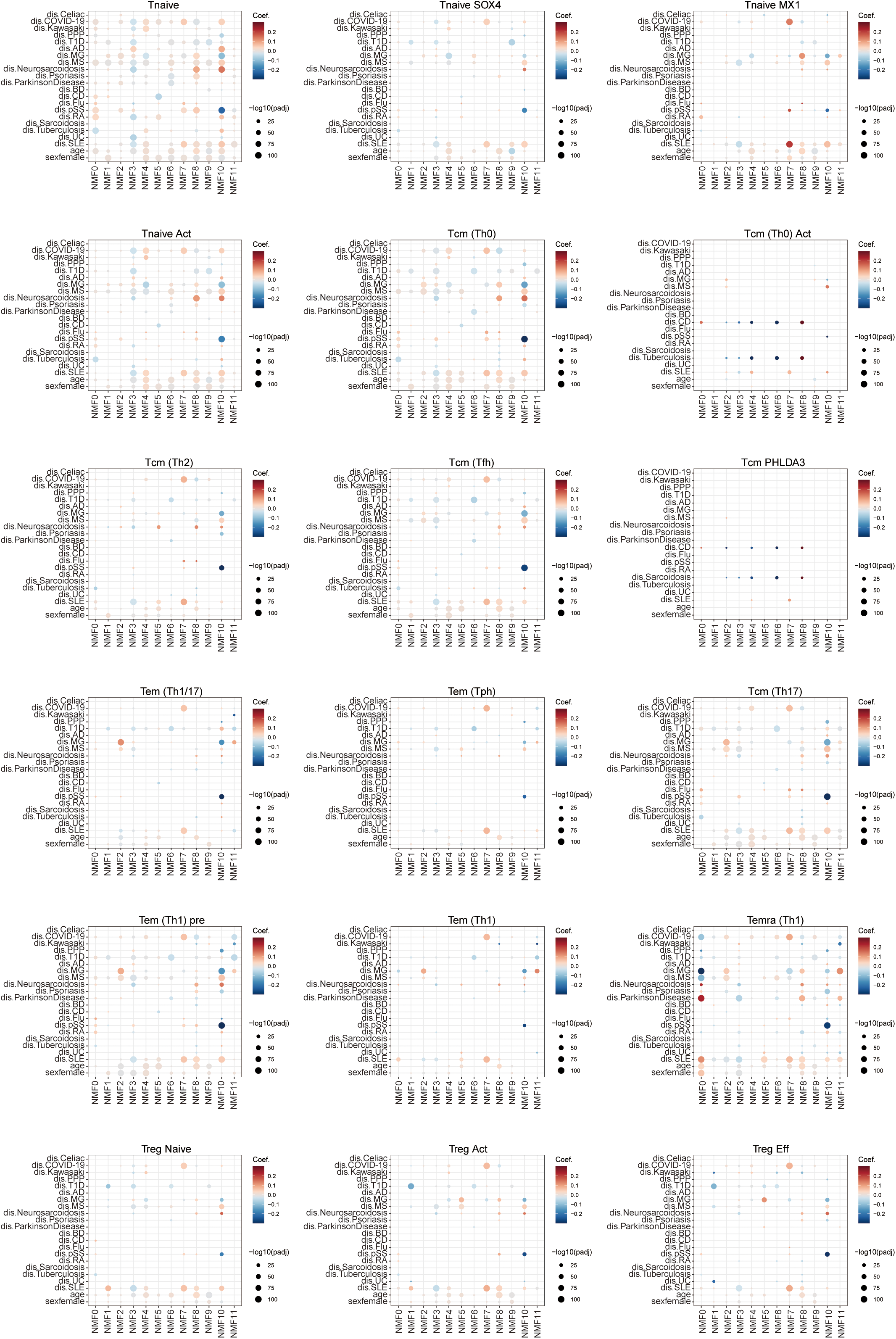
NMF cell feature changes depending on diseases. Dot plots depicting NMF cell feature changes in each cell type. Dot colors show coefficients, and sizes show the significance of GLM. GLM was performed with a model, NMF cell feature ∼ disease + age + gender + project. Only significant dots (p_adj_ < 0.05) are shown.

**Figure S10.**
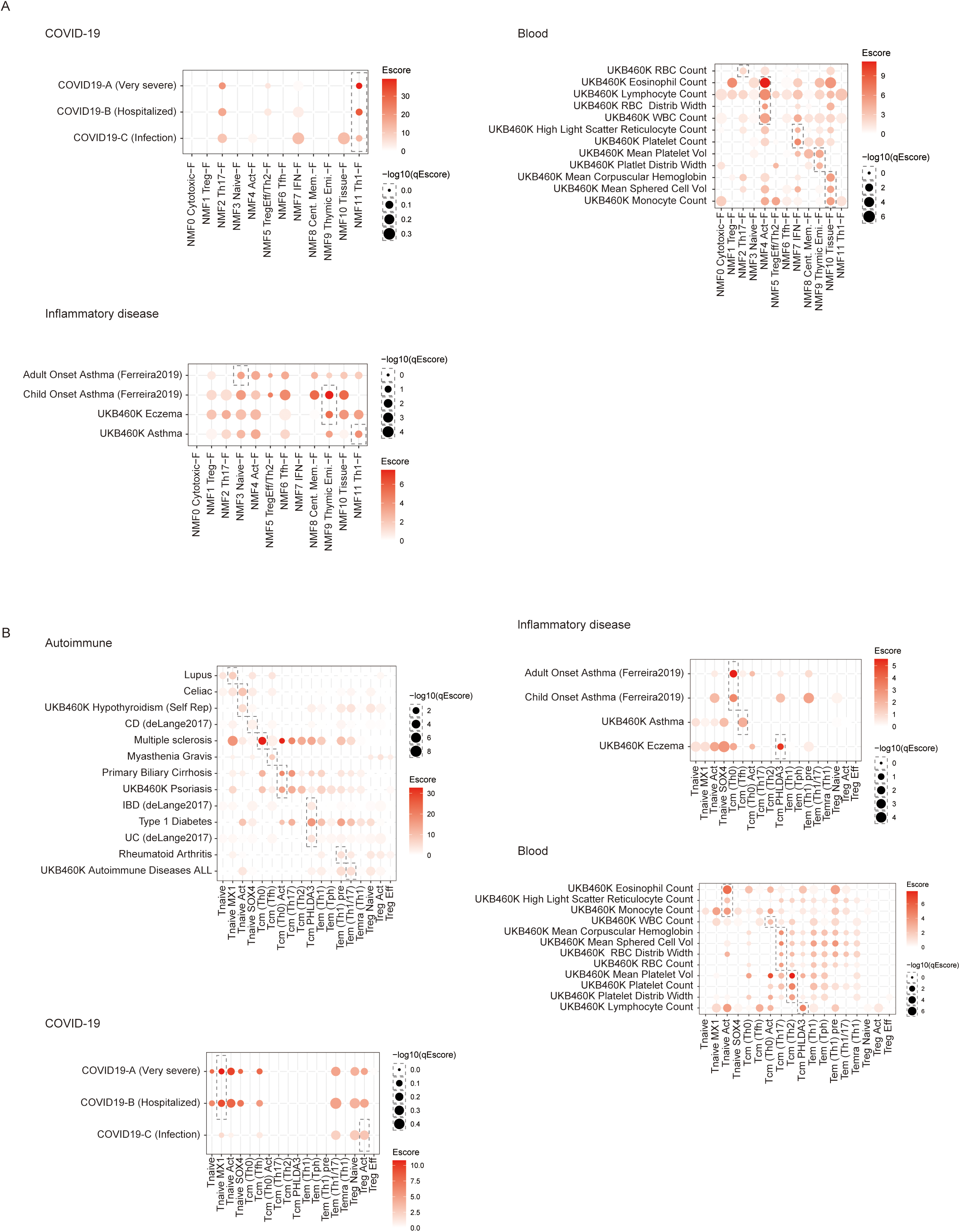
Partitioned heritability. (A and B) Dot plots showing partitioned heritability of diseases across NMF gene features (A) or cell types (B). Duplicated traits were removed for the visualization. Full statistics are shown in Tables S11 and 12.

## Supplementary Note - The applications of NMFproj -

Miyara classification (bulk RNA-seq)

We utilized NMFproj to bulk RNA-seq data of sorted peripheral CD4^+^ T cells for each fraction in the Miyara classification ^30^, which classifies CD4^+^ T cells by CD25 and CD45RA (Figure S3E, POH: 0.542). Consistent with previous findings ^30^, Fr. I and Fr. II exhibited high NMF1 (Treg-F), while Fr. V and Fr. IV showed low NMF1 (Treg-F). Similarly, NMF3 (Naive-F) was found to be high in Fr. I and Fr. VI, and low in Fr. II and Fr. V, which also aligns with existing knowledge. Furthermore, Fr. III has been reported to possess weak suppressive activity and a Th17-like phenotype ^30^, and this was concordant with our observation that NMF1 (Treg-F) was lower in Fr. III compared to Fr. I and Fr. II, and NMF2 (Th17-F) was higher.

Tph cells (bulk RNA-seq)

Profiling of sorted Tph cells also revealed that Tph is a population with both Tfhness (NMF6) and Th1ness (NMF11) (Figure S3F, POH: 0.134).

iTreg cells (bulk RNA-seq)

When we applied NMFproj to *in vitro* induced Tregs (iTregs) ^31^, NMF1 (Treg-F) was higher in nTreg cells than iTreg cells (Figure S3G). iTreg cells made in conditions to stabilize Treg function with CD28 depletion and two times resting showed higher NMF1 (Treg-F) than other iTreg cells concordantly with the experimentally measured suppressive functions. This suggests that NMFproj can be used for the evaluation of Tregness in a genome-wide manner rather than the tracing of single or a few genes, as performed in most studies, as well as monitoring of unwanted polarization.

Pan-cancer CD4^+^ T cells (scRNA-seq)

We analyzed scRNA-seq of the pan-cancer CD4^+^ T cell dataset ^15^ by utilizing NMFproj (Figure S4A; POH: 0.53). The 12 factors were also conserved in the tumor microenvironments. Most Treg cells possessed high NMF5 (TregEff/Th2-F), indicating Treg activation in tumor environments. NMF10 (Tissue-F) was broadly high in tumor CD4^+^ T cells.

Mouse splenocytes (scRNA-seq)

Single-cell data from mouse splenocytes ^33^ were analyzed to confirm whether cross-species projection was possible. We found that NMFproj could capture not only relatively large populations of Treg and Tfh but also small populations such as Th17 and Th2, which were not indicated in the original paper (Figure S4B, POH: 0.394). This result indicates that cross-species projection is also possible and, moreover, that NMFproj is informative even for single-cell data with a small number of cells.

Cross-tissue immune cells (scRNA-seq)

We reanalyzed the single-cell cross-tissue dataset ^32^ to investigate how the gene features created using peripheral blood would behave in organs (Figure S5). The projected cell features were concordant with the defined cell population, except for the absence of cytotoxic CD4^+^ Temra. Then, we noticed that the cytotoxic CD4^+^ Temra was incorrectly defined as a CD8^+^ T population and part of CD8^+^ T as CD4^+^ T in the original report. We had similar experiences where CD4^+^ T cells and CD8^+^ T cells were not well separated and embedded as a mixed cluster in single-cell analysis. We hypothesized that CD4^+^ T and CD8^+^ T use similar genetic programs, examined the POH of each cell type, and found that, surprisingly, CD8^+^ T cells, innate T cells, and even B cells and myeloid cells marked relatively high POH. When Evar was examined, CD8^+^ T cells and innate T cells were found to preferentially use NMF0 (cytotoxic-F), while B cells and Myeloid cells used NMF5 (TregEff/Th2-F). These observations suggested that gene programs were evolutionally developed and conserved across cell populations. Examination of POH in CD4^+^ T cells by tissue showed that POH was high in peripheral blood and secondary lymphoid tissues, while POH was low in tissues such as the liver and muscle, suggesting that the gene features defined using peripheral CD4^+^ T cells do not fully represent the tissue response. In addition, the Evar of TregEff/Th2-F, Th17-F, and Th1-F were found to be high in tissues, suggesting that polarization is a prominent event in tissues.

Sorted CD4^+^ T cell fractions from autoimmune patients (bulk RNA-seq)

NMFproj was adapted to the ImmuNexUT dataset ^35^, which contains sorted CD4^+^ T cell fractions across autoimmune diseases to capture qualitative changes in each CD4^+^ T cell population (Figure S6 POH: 0.586). The most prominent variation is NMF7 (IFN-F), which is elevated across cell types in SLE and MCTD. Also, Tregness (NMF1) decreased in SLE in naive Treg, indicating Treg dysfunction in SLE patients.

## Notes

### Competing Interest Statement

The authors have declared no competing interest.

